# Selenite transport by SLC26A6 contributes to redox homeostasis and ferroptosis resistance

**DOI:** 10.64898/2026.11.16.767030

**Authors:** Sena Atici, Panagiotis Kreouzis, Gholamreza Fazeli, Maria Schwarz, Zhiyi Chen, David Tippett, Ancély Ferreira dos Santos, Ana Clara Najenson, Mami Sato, Shibo Sun, Florencio Porto Freitas, Jonas Gröninger, Kamini Kaushal, Mostafa Ejtehadifar, Hana Nuskova, Nikolas Gunkel, Felix Knauf, Namgyu Lee, Enrico Girardi, Giulio Superti-Furga, Martin Fassnacht, Chen Deng, Eikan Mishima, Bettina Proneth, Marcus Conrad, Hamed Alborzinia, Anna Patricia Kipp, Raimund Dutzler, José Pedro Friedmann Angeli

**Affiliations:** Rudolf Virchow Zentrum (RVZ), Center for Integrative and Translational Bioimaging, University of Wuerzburg, Wuerzburg, Germany; Department of Biochemistry, University of Zurich, Zurich, Switzerland; Department of Nutritional Physiology, Institute of Nutritional Sciences, Friedrich Schiller University Jena, Jena, Germany; Department of Nephrology and Medical Intensive Care, Charité–Universitätsmedizin Berlin, Berlin, Germany; Heidelberg Institute for Stem Cell Technology and Experimental Medicine (HI-STEM gGmbH), Heidelberg, Germany; Division of Stem Cells and Cancer, German Cancer Research Center (DKFZ), Heidelberg, Germany; Research Group Cancer Drug Development, German Cancer Research Center (DKFZ), Heidelberg, Germany; Department of Biomedical Science & Engineering, Dankook University, Cheonan, Republic of Korea; CeMM Research Center for Molecular Medicine of the Austrian Academy of Sciences, Vienna, Austria; Solgate GmbH, Klosterneuburg, Austria; Medical University of Vienna, Center for Physiology and Pharmacology, Vienna, Austria; Division of Endocrinology and Diabetes, Department of Medicine, University Hospital, University of Würzburg, Würzburg, Germany; Helmholtz Zentrum München, Institute of Metabolism and Cell Death, Molecular Targets and Therapeutics Center, Neuherberg, Germany; Translational Redox Biology, Technical University of Munich (TUM), TUM Natural School of Sciences, Garching, Germany; Comprehensive Cancer Center Mainfranken, University hospital Würzburg, Würzburg, Germany

**Keywords:** Selenium, Selenoproteins, Solute Carriers, SLC26A6, CRISPR screen, Selenite

## Abstract

**Highlights:**

– SLC26A6 promotes selenite import.
– SLC26A6-mediated selenite import promotes selenoprotein biosynthesis.
– SLC26A6 protects cells from ferroptosis in a selenite dependent manner.
– The small molecule A6i inhibits selenium uptake by SLC26A6, restoring ferroptosis sensitivity.

Graphical Abstract

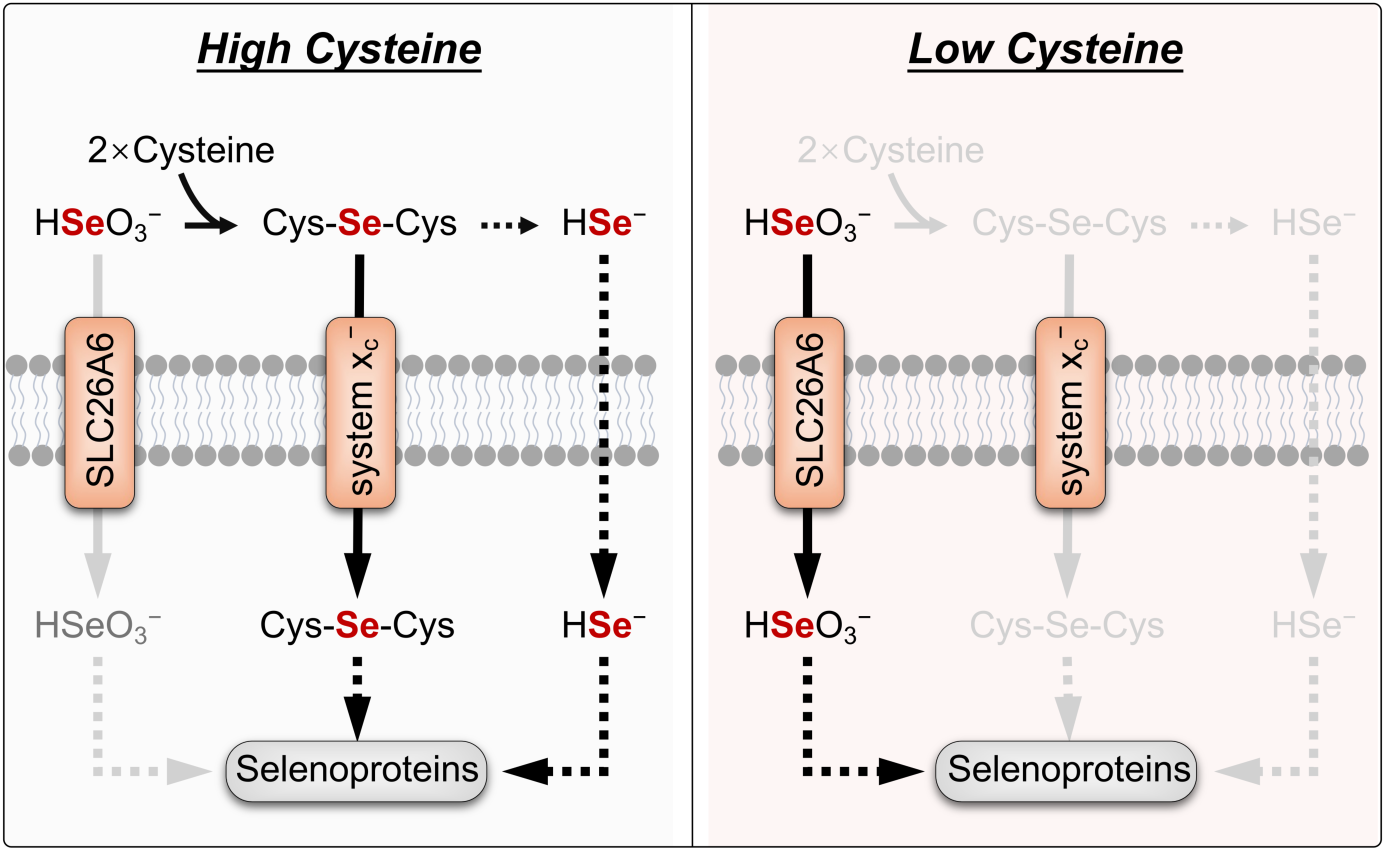

Selenium is an essential micronutrient widely found in dietary supplements and fortified foods and is required for selenoprotein synthesis and function. Although the molecular mechanisms of selenium-dependent selenoprotein translation are well understood, its uptake mechanisms remain only partially characterized. Using a CRISPR-based loss-of-function screen, we identify the solute carrier SLC26A6 as a direct selenite transporter. Using biochemical, cellular, and *in vivo* studies, we show that SLC26A6 directly transports selenite and promotes selenoprotein translation, redox homeostasis, and ferroptosis resistance. Additionally, we report the first human-specific SLC26A6 inhibitor and determine the structure of the SLC26A6-inhibitor complex by cryo-electron microscopy. These findings identify a previously uncharacterized route of inorganic selenium uptake and establish SLC26A6 as a pharmacologically tractable determinant of selenoprotein synthesis, redox homeostasis, and ferroptosis resistance.

## Introduction

Selenium is an essential micronutrient required for the biosynthesis of selenocysteine, the rare amino acid defining a small class of proteins known as selenoproteins ^1–3^. Selenoproteins are key regulators of redox homeostasis, including glutathione peroxidase 4 (GPX4), a unique enzyme required to suppress lipid peroxidation and ferroptotic cell death ^4,5^. Selenium availability and uptake therefore directly link this micronutrient to the function of these proteins.

Dietary selenium includes organic forms (i.e selenomethionine and selenocysteine), and inorganic forms (i.e, selenite and selenate) ^6,7^. Cells acquire selenium through multiple routes, including LRP8, which mediates uptake of selenoprotein P ^8^, or indirectly through system x_c_^−^. SLC4A1 has been described as a selenite transporter and is exclusively present in erythrocytes ^9^, but a broadly expressed mechanism for direct inorganic selenium uptake remains undefined. Selenite (SeO_3_^2−^) is widely used in cell culture and as a nutritional supplement in humans to support selenoprotein synthesis ^10^, its cellular uptake is thought to involve, at least in part, a thiol-mediated extracellular reduction of selenite and subsequent uptake through system x_c_^− 11,12^. Whether mammalian cells possess a direct, thiol-independent selenite transporter has remained unresolved. Identifying such a pathway would establish a fundamental component of selenium biology and reveal a potential point of control over selenium-dependent processes, including ferroptosis.

Here, we report on the identification of the solute carrier 26 family member A6 (SLC26A6), a member of a large family of anion transporters ^13^, as a direct selenite transporter that supports selenoprotein synthesis and ferroptosis resistance. Genetic and functional analyses *in vitro* and *in vivo* demonstrate that SLC26A6 promotes selenite uptake and maintains selenoprotein production, particularly under physiological and low extracellular thiol concentrations. We further identify A6i, a selective small-molecule inhibitor of human SLC26A6, and show that pharmacological blockade of the transporter reduces selenoprotein production and sensitizes cells to ferroptosis.

## Results

### Identification of SLC26A6 as a selenite transporter

To identify transporters mediating selenite uptake, we performed a CRISPR knockout screen targeting solute carrier (SLC) genes in A549 cells, which are sensitive to selenite ^3,11^ (Figure 1A). Given that selenite is cytotoxic to cells at high concentrations ^14^, the loss of a functional uptake pathway was expected to confer a survival advantage and allow the identification of a putative transporter. The screen identified SLC7A11, a component of system x_c_^−^ previously implicated in indirect selenium uptake ^11,12^, and the anion transporter SLC26A6 (Figure 1B). Interestingly, SLC26A6 has been reported to transport bicarbonate ^13,15^, an anion structurally similar to selenite, suggesting it could act as a direct transporter for selenite.

**Figure 1:**
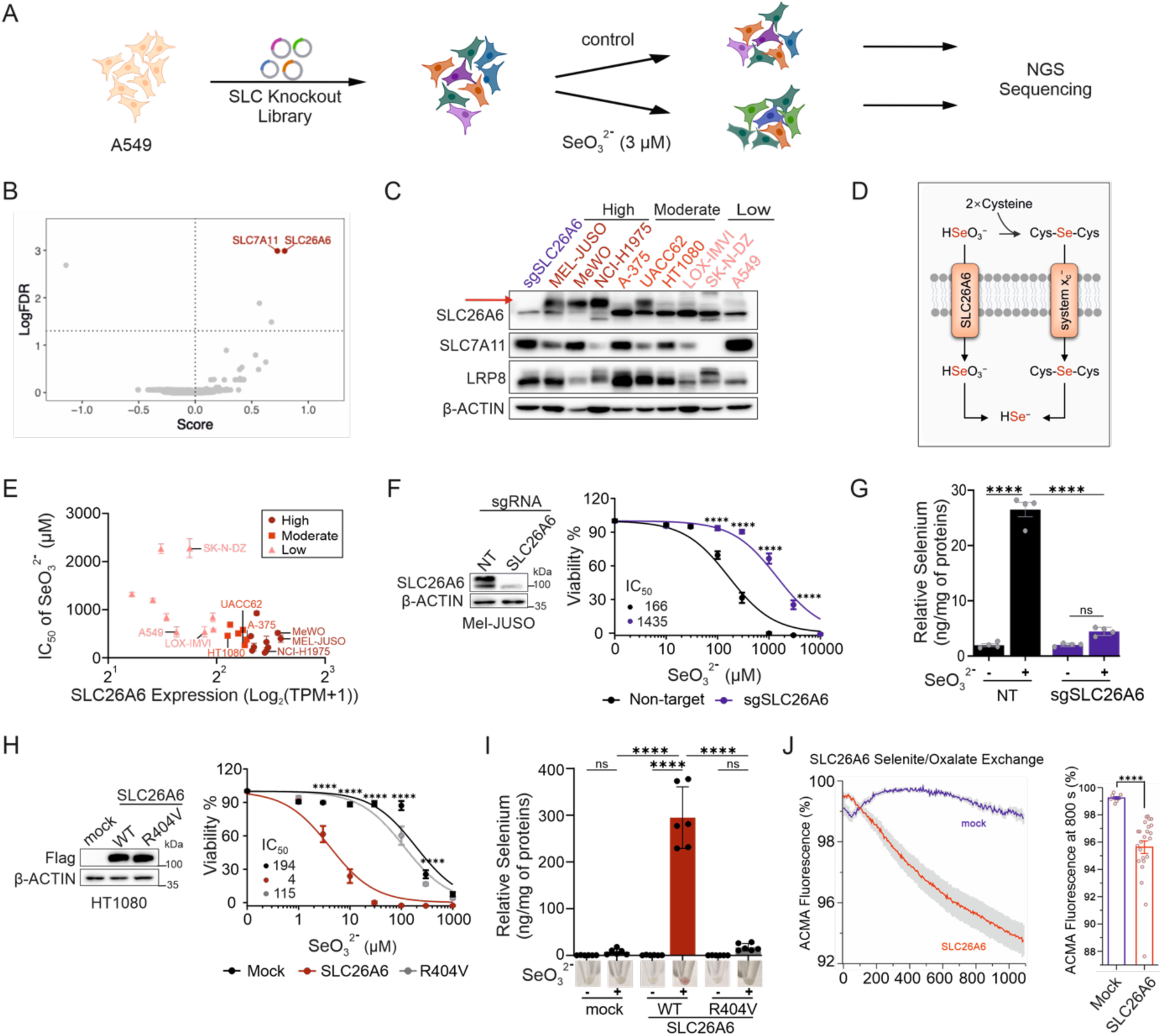
SLC26A6-mediated selenite uptake promotes selenite toxicity. **(A)** Schematic of a CRISPR knock-out screen focused on solute carrier (SLC) genes. **(B)** LogFDR results for CRISPR knock-out screen in A549 lung carcinoma cells upon supplementation with 3µM of selenite, showing SLC7A11 and SLC26A6 as top hits. **(C)** Immunoblot analysis of SLC26A6, SLC7A11 and LRP8 from different cancer cell line. Cell lines were grouped as high, moderate and low based on their SLC26A6 protein levels. **(D)** Schematic representation of the proposed pathways for importing inorganic selenium. **(E)** Correlation of the IC_50_ values for selenite in different cancer cell lines and their SLC26A6 expression levels. Cells were grouped according to their SLC26A6 expression levels. Expression values are derived from DepMap (https://depmap.org). Pearson correlation: *r* =-0.68. **(F)** Immunoblot analysis and viability of Mel-JUSO cells transduced with non-target or SLC26A6 sgRNAs, in the presence of increasing concentration of sodium selenite. Data are mean ± SEM of 3 wells of 96-well plate from 3 independent experiments. Data are normalized to vehicle control of each cell line. **** p<0.0001. Statistical analysis was performed using a two-way ANOVA with adjustment for multiple comparisons. **(G)** Total selenium content of Mel-JUSO cells transduced with non-target or SLC26A6 sgRNAs after treatment with 50 μM of selenite for 6 hours. Total selenium was measured using TXRF method. 1mg/L Yttrium was used as standard. Measurements are normalized to protein levels. **** p<0.0001. Statistical analysis was performed using a two-way ANOVA with adjustment for multiple comparisons. **(H)** Immunoblot analysis of HT1080 cells transduced with mock, FLAG-tagged WT or FLAG-tagged R404V mutant of SLC26A6. The right panel shows the viability of cells in the presence of increasing concentrations of selenite. Data are mean ± SEM of 3 wells of 96-well plate from 3 independent experiments. Data are normalized to the vehicle control of each cell line. **** p<0.0001. Statistical analysis was performed using a two-way ANOVA with adjustment for multiple comparisons. **(I)** Total selenium content of HT1080 mock, SLC26A6 WT and R404V overexpressing cells after treatment with 50μM of selenite for 6 hours. **** p<0.0001. Statistical analysis was performed using a two-way ANOVA with adjustment for multiple comparisons. Representative images of the tubes after 50µM selenite treatment are shown under each treatment. Reddish color indicates the presence of elemental selenium. **(J)** Electrogenic oxalate/selenite exchange into proteoliposomes containing reconstituted SLC26A6 in comparison to mock liposomes. Transport is assayed via the parallel import of H^+^ with the pH-sensitive fluorophore ACMA. Left, time dependence of ion transport. Traces show means of 24 replicates from 2 independent reconstitutions for SLC26A6 containing proteoliposomes and 8 replicates from 2 independent reconstitutions for mock. Right, comparison of the relative fluorescence decrease after 800 Sec. Individual values are indicated. **** p<0.0001. Statistical analysis was performed using a Welch’s ANOVA.

To test the relative contribution of putative selenite transporters in different cancer cell lines, we analyzed the protein levels of SLC26A6, SLC7A11, and LRP8 (Figure 1C). Given the previously reported contribution of system x_c_^−^ to selenite uptake, likely via the formation of mixed cysteine-selenite intermediates (Figure 1D), and to isolate the contribution of SLC26A6, subsequent experiments were performed under cystine-restricted conditions, unless stated otherwise. Using the DepMap expression datasss and experimentally determined IC_50_ values for selenite across cell lines, we found that SLC26A6 expression negatively correlated with selenite IC_50_ values (*r* = −0.68; Figure 1E). These findings implicated SLC26A6 in mediating selenite toxicity and suggested that selenite-induced cell death is a suitable proxy for studying this transporter’s activity in cells.

We tested this relationship by deleting SLC26A6 in A549 cells (Figures S1A and S1B) and in cell lines with high endogenous expression (Mel-JUSO, Me-Wo, and NCI-H1975; Figures 1F, S1C and S1D). SLC26A6 loss conferred resistance to selenite under cystine-restricted conditions. Conversely, overexpression of wild-type SLC26A6 in HT1080 cells increased selenite sensitivity, whereas the bicarbonate transport-inactive mutant R404V ^16^ failed to sensitize the cells to selenite (Figure 1H), an effect recapitulated in a larger panel of cells (Figure S1E). Erastin-mediated inhibition of SLC7A11 did not substantially attenuate this effect (Figure S1F), supporting a role for SLC26A6 in thiol-independent selenite uptake. Moreover, SLC26A6 overexpression did not increase sensitivity to selenate or selenocystine (Figure S1G), showing the selectivity of this transporter for selenite.

To directly assess selenium uptake, we measured intracellular selenium by total reflection x-ray fluorescence (TXRF) upon selenite exposure. Mel-JUSO or NCI-H1975 cells with high endogenous SLC26A6 expression accumulated selenium, whereas SLC26A6-knockout cells did not show a significant increase (Figures 1G and S1H). Similarly, overexpression of wild-type SLC26A6, but not the transport-inactive R404V mutant, increased selenium accumulation in HT1080 cells (Cys-free medium, Figure 1I and Cys-containing medium, Figure S1I). Selenite-treated cell pellets also developed a reddish color consistent with elemental selenium accumulation ^17^ (Figure 1I). This effect was specific to selenite and was not observed with other selenium sources (Figure S1J). These findings support a role for SLC26A6 in cellular selenite uptake.

To further assess SLC26A6 specificity in selenite transport, we compared the effects of overexpressing other SLC26 and SLC4 family members implicated in bicarbonate transport, including SLC4A1, which has been reported to contribute to selenite transport in erythrocytes^9^ ^9^. Unlike SLC26A6, no selenite sensitivity or intracellular selenium accumulation was observed for the other transporters tested, except for the previously reported erythrocyte SLC4A1 (Figures S2 and S3). To further confirm this in an unbiased manner, we performed an independent CRISPR activation screen targeting 388 solute carriers. This screen, designed to identify transporters whose increased expression promotes selenite-mediated toxicity, independently identified SLC26A6 and SLC7A11 as sensitizers (Figure S3L), providing orthogonal validation of SLC26A6’s relative contribution as a selenite transporter (note that the library used for the screen did not include SLC4A1-targeting sgRNAs) ^18^.

To test whether SLC26A6 directly transports selenite, we reconstituted purified protein produced in HEK293 cells into liposomes. In proteoliposomes containing selenite, adding external oxalate induced an ACMA-detectable response, resulting from parallel proton movement accompanying the electrogenic anion exchange of divalent oxalate and monovalent HSeO_3_^−^ (Figure 1J). In contrast, no comparable response was observed with the impermeant anion gluconate trapped inside the liposomes (Figures S4A and S4B). Furthermore, a cryo-EM structure of SLC26A6 prepared in gluconate-containing conditions and exposed to selenite before vitrification (Figures S4C to S4H, and Table S3) revealed a 3.65-Å map of an inward-facing conformation, similar to the previously reported chloride-bound structure ^16^, with weak residual density in the substrate-binding site consistent with bound selenite, the only transported anion present during sample preparation (Figure S4H).

### SLC26A6 supports selenoprotein biosynthesis and redox homeostasis

Having established that SLC26A6 promotes selenite import, we next asked whether this transport activity could support selenoprotein biosynthesis at the low nanomolar selenite concentrations typically used in supplementation experiments. In line with the essential role of the selenoprotein GPX4 in suppressing ferroptosis ^19,20^, SLC26A6 overexpression protected cells against GPX4 inhibitors (FIN56 and RSL3) (Figure 2A).

**Figure 2:**
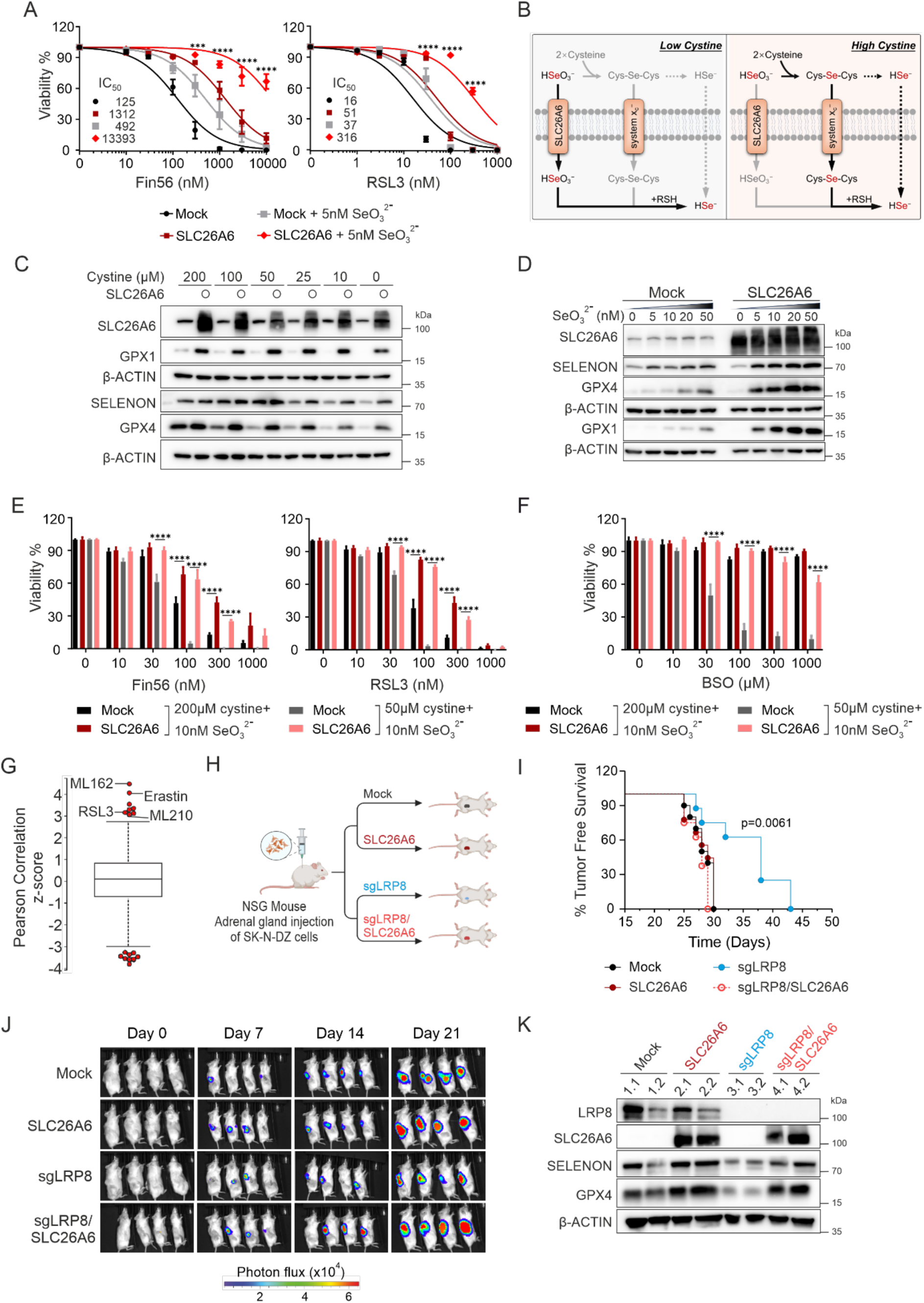
SLC26A6-mediated selenite uptake enhances selenoprotein expression and ferroptosis resistance. **(A)** Viability of mock or SLC26A6 overexpressing cells in the presence of increasing concentration of Fin56 (left) or RSL3 (right) in combination with selenite. Data are mean ± SEM of 3 wells of 96-well plate from 2 independent experiments. Data are normalized to vehicle control of each cell line. Cells were pre-incubated for 2 days with selenite prior to Fin56 or RSL3 treatment under cystine containing conditions. *** p<0.001, **** p<0.0001. Statistical analysis was performed using a two-way ANOVA with adjustment for multiple comparisons. **(B)** A scheme illustrating the change in relative contributions of SLC26A6 and system x_c_^−^ to selenoprotein production through selenite uptake. Contribution of system x_c_^−^ dominates at high cystine concentrations, while SLC26A6’ role becomes more apparent at low physiological cystine concentrations. **(C)** Immunoblot analysis of the selenoproteins GPX4, GPX1, and SELENON in response to 10 nM of selenite, 500nM Lip1, and indicated concentrations of cystine in mock or SLC26A6 overexpressing HT1080 cells. **(D)** Immunoblot analysis of the selenoproteins GPX4, GPX1, and SELENON in response to increasing nanomolar concentrations of selenite and 500nM Lip1 in mock or SLC26A6 overexpressing HT1080 cells in cystine-free condition. **(E)** Viability of mock or SLC26A6 overexpressing cells in the presence of increasing concentration of Fin56 (left) or RSL3 (right) in combination with 10 nM selenite. Data are mean ± SEM of 2 wells of 96-well plate from 2 independent experiments. Data are normalized to vehicle control of each cell line. Cells were pre-incubated for 2 days with selenite and the indicated cystine concentrations prior to Fin56 or RSL3 treatment. **** p<0.0001. Statistical analysis was performed using a two-way ANOVA with adjustment for multiple comparisons. **(F)** Viability of mock or SLC26A6 overexpressing cells in the presence of increasing concentration of BSO in combination with 10 nM selenite. Data are mean ± SEM of 2 wells of 96-well plate from 2 experiments. Data are normalized to vehicle control of each cell line. Cells were pre-incubated for 2 days with the indicated cystine concentrations in combination with selenite prior to BSO treatment. **** p<0.0001. Statistical analysis was performed using a two-way ANOVA with adjustment for multiple comparisons. **(G)** High expression of SLC26A6 correlates with resistance to GPX4 inhibitors. Drug sensitivity correlation with SLC26A6 gene expression levels was retrieved from Cancer Therapeutics Response Portal (CTRPv2.1, https://portals.broadinstitute.org/ctrp.v2.1). **(H)** Schematic representation of the orthotopic implantation of WT, LRP8-deficient (sgLRP8), SLC26A6 overexpressing (SLC26A6), and LRP8-deficient with SLC26A6 overexpressing (sgLRP8/SLC26A6) neuroblastoma SK-N-DZ cells in adrenal gland of mice. **(I)** Tumor free survival of mice orthotopically implanted with WT (n=10), LRP8-deficient (sgLRP8, n=8), SLC26A6 overexpressing (SLC26A6, n=9), or LRP8-deficient with SLC26A6 overexpressing (sgLRP8/SLC26A6, n=8) SK-N-DZ cells. Statistical analysis was performed using a Mantel-Cox test. **(J)** Representative bioluminescence images of tumor formation for mice with implanted cells shown in (H) and (I). Images are shown on a linear color scale (photons/sec/cm^2^/sr). **(K)** Immunoblot analysis of SLC26A6, LRP8, SELENON and GPX4 from the orthotopic tumor samples collected after sacrificing animals.

Importantly, our genetic screen suggests that SLC26A6 and system x_c_^−^ provide parallel routes for selenium uptake, possibly with their relative contributions depending on extracellular cysteine availability. Cysteine was shown to react directly with selenite to form selenodicysteine (Cys–Se–Cys), a selenotrisulfide proposed to be a substrate of system x_c_^− 12,21^. Thus, extracellular cysteine levels could determine the preferred route for uptake (Figure 2B).

To test this model, we varied extracellular cysteine/cystine concentrations and assessed selenoprotein expression in the presence of 10 nM selenite. At high cysteine concentrations (200 μM), SLC26A6 had little effect on selenoprotein expression, with an effect detected only for GPX1, a selenoprotein known to be more sensitive to selenium availability ^22^. By contrast, at 50 μM cysteine (typical plasma concentrations) and under cysteine-restricted conditions, SLC26A6 was the main entry point for selenite and, consequently, selenoprotein production (Figures 2C and S5A). These findings are consistent with a shift from selenium uptake through a cysteine-derived intermediate at high cysteine availability toward direct selenite uptake through SLC26A6 as cysteine becomes limiting. Further supporting this model, increasing selenite concentrations under cysteine-restricted conditions enhanced selenoprotein expression in cells expressing wild-type SLC26A6, but not the R404V mutant (Figures 2D and S5B). Conversely, SLC26A6 deletion reduced selenoprotein levels and prevented their increase following selenite supplementation under cysteine-free conditions (Figure S5C). This effect also depended on the selenium source, as selenate did not alter selenoprotein levels in either control or SLC26A6-overexpressing cells, whereas selenocystine increased these levels similarly in both groups (Figures S5D to S5G). Together, these results support a specific role for SLC26A6 in selenite-promoted selenoprotein biosynthesis, particularly at physiological concentrations and at low extracellular cysteine levels.

We next asked whether SLC26A6-mediated enhanced selenoprotein production promotes redox homeostasis and, more specifically, resistance to ferroptosis. At physiological extracellular cysteine concentrations, SLC26A6 overexpression protected cells against several paradigms of ferroptosis and oxidative stress induction, including GPX4 inhibitors (FIN56 and RSL3), and glutathione depletion (BSO) (Figures 2E and F). This protection persisted during SLC7A11 inhibition with erastin, which also failed to abolish the increase in selenoprotein expression driven by SLC26A6 (Figures S6A and S6B). Consistent with these findings, low-dose selenite reduced lipid peroxidation induced by FIN56, as assessed by the oxidized-to-reduced C11-BODIPY fluorescence ratio, in the presence of erastin (Figure S6C). Cysteine restriction further increased the dependence of selenoprotein synthesis on SLC26A6 (Figures S6D and S6E). SLC26A6 also protected cells against cysteine-withdrawal-induced ferroptosis in a selenite-dependent manner (Figure S6F). Conversely, SLC26A6 deletion sensitized Mel-JUSO cells to cell death, which was rescued by liproxstatin-1 (Lip-1) (Figure S6G). Finally, genetic deletion of GPX4 abolished the protection conferred by SLC26A6 overexpression (Figures S6H and S6I), establishing GPX4 as a necessary mediator of this effect. Further support for SLC26A6’s role in protecting cells from oxidative stress and ferroptosis comes from the Cancer Therapeutics Response Portal, which reported an association between SLC26A6 expression and resistance to GPX4 inhibitors across cancer cell lines (Figure 2G). Together, these findings establish SLC26A6-mediated selenite uptake as a determinant of selenoprotein biosynthesis and ferroptosis resistance.

To challenge the relevance of SLC26A6-mediated selenite uptake *in vivo*, we used an orthotopic neuroblastoma xenograft model in which MYCN-amplified tumors depend on the SELENOP receptor LRP8 for survival ^23–25^. SK-N-DZ cells express low levels of both SLC7A11 and SLC26A6 (Figure 1C), potentially contributing to this dependency. SLC26A6 overexpression restored selenite-dependent GPX4 translation in LRP8-knockout (KO) cells (Figure S6J). Consistent with this, selenite supplementation selectively improved the survival of SLC26A6-overexpressing LRP8-KO cells, whereas liproxstatin-1 (Lip-1) rescued cells irrespective of SLC26A6 expression (Figure S6K). Thus, SLC26A6-mediated selenite uptake can compensate for impaired LRP8-dependent selenium acquisition. We next implanted SK-N-DZ cells into the adrenal glands of mice (Figure 2H). LRP8 deletion delayed tumor onset, as previously reported ^23^, whereas SLC26A6 overexpression reversed this effect (Figures 2I and 2J). Tumors derived from SLC26A6-overexpressing LRP8-KO cells also showed increased selenoprotein expression, demonstrated here for GPX4 and SELENON (Figure 2K). These findings show that SLC26A6-mediated selenium uptake can compensate for LRP8 loss, sustaining selenoprotein expression and tumor growth *in vitro* and *in vivo*.

### Identification and characterization of an SLC26A6 inhibitor

We reasoned that pharmacological inhibition of SLC26A6-mediated selenite uptake could be a potential tool to sensitize cells to ferroptosis. Because sulfite resembles selenite structurally, we first tested whether high concentrations of sulfite could inhibit selenite uptake. Accordingly, sulfite, but not sulfate, rescued SLC26A6-overexpressing cells from selenite toxicity (Figure S7A), providing experimental evidence that an SLC26A6 inhibitor could protect cells against selenite-induced toxicity. We therefore screened a ∼30,000-compound library using selenite-induced toxicity as a cellular readout of SLC26A6 activity (Figure 3A). Upon validation, the screening campaign identified a single compound with robust inhibitory activity in our cellular assay, which we termed A6i (Figure 3B). A6i rescued cells from selenite toxicity with an apparent EC_50_ of 2-3 μM (Figure 3C). Structure–activity studies of 14 related compounds identified a hydroxyl group on the benzene ring as essential for inhibitor activity. Consistent with this finding, A6i.05, in which the hydroxyl group was replaced by sulfur (Figure 3B), failed to show inhibitory activity (Figure 3C and Table S1).

**Figure 3:**
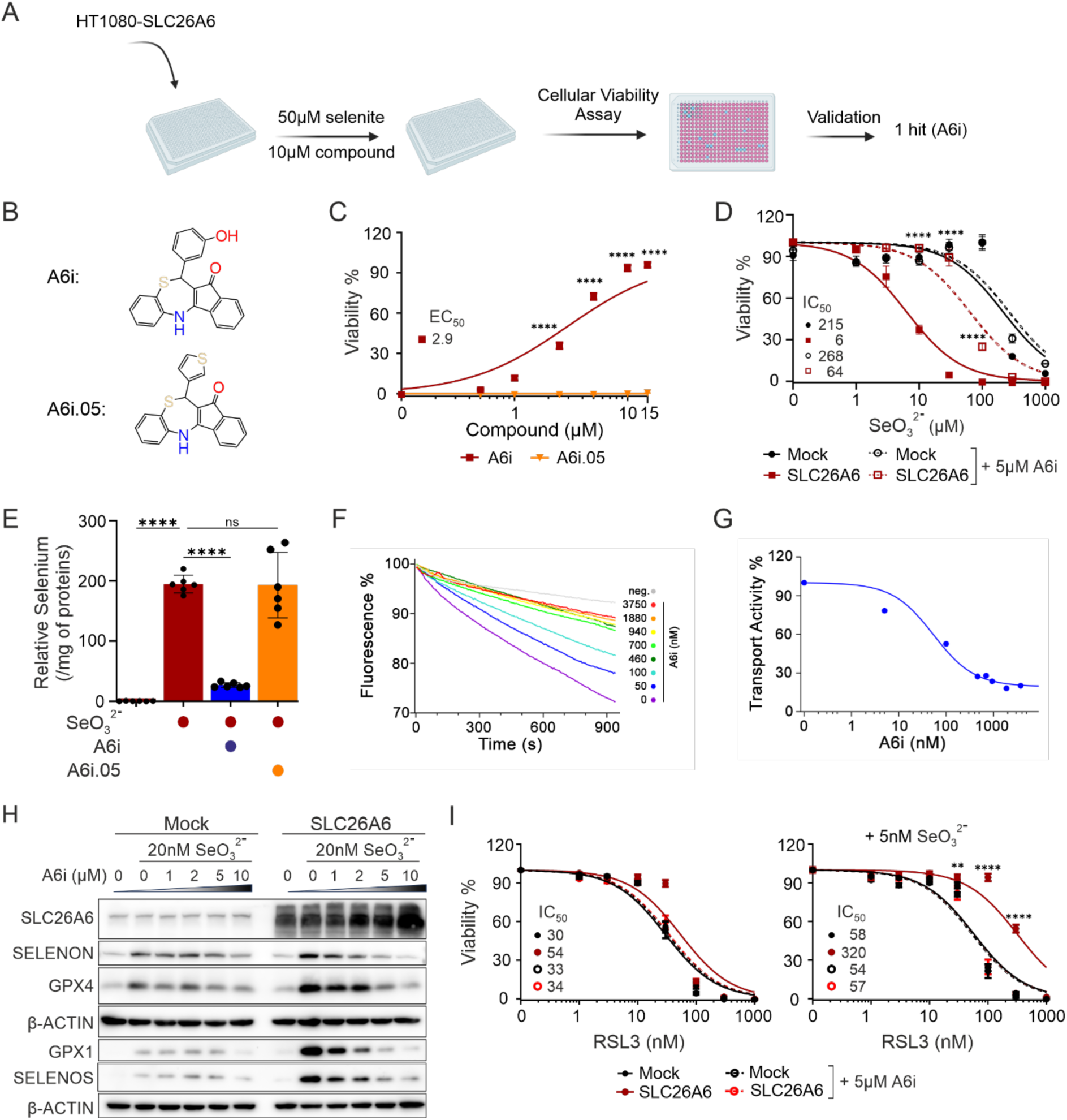
Selenite toxicity-based phenotypic screen identifies SLC26A6 inhibitors. **(A)** A scheme illustrating a small-molecule library screen on SLC26A6 overexpressing cells. **(B)** Structures of A6i and its inactive analog A6i.05. **(C)** Viability of SLC26A6 overexpressing cells treated with 50 μM of sodium selenite together with the increasing concentration of A6i or A6i inactive analog A6i.05 in the absence of Lip1. Data are mean ± SEM of 3 wells of 96-well plate from 3 independent experiments. Data are normalized to 500nM Lip1 containing control of each cell line. **(D)** Viability of mock and SLC26A6 overexpressing cells, in the presence of increasing concentration of selenite, with or without 5µM of A6i. Data are mean ± SEM of 3 wells of 96-well plate from 3 independent experiments. Data are normalized to the vehicle control of each cell line. **** p<0.0001. Statistical analysis was performed using a two-way ANOVA with adjustment for multiple comparisons. SLC26A6 overexpressing cells with A6i are compared with those without A6i. **(E)** Total selenium content of SLC26A6 overexpressing cells after treatment with 50 μM of selenite in combination with A6i or its inactive analog A6i.05. **** p<0.0001. Statistical analysis was performed using a two-way ANOVA with adjustment for multiple comparisons. **(F)** Concentration-dependent inhibition of oxalate/Cl**^−^** exchange into proteoliposomes containing reconstituted SLC26A6 by A6i. Transport is assayed via the parallel import of H^+^ with the pH sensitive fluorophore ACMA. Traces show means of five replicates from 2 independent reconstitutions. **(G)** Concentration-response relationship of inhibition of SLC26A6 by A6i. Transport activities were obtained from (F). The fit shows a binding isotherm with an IC_50_ of 54 nM. **(H)** Immunoblot analysis of the selenoproteins GPX4, GPX1, SELENON, and SELENOS in response to increasing concentrations of A6i with or without 20 nM selenite in mock or SLC26A6 overexpressing HT1080 cells. **(I)** Viability of mock and SLC26A6 overexpressing cells in the presence of increasing concentration of RSL3 with or without 5 µM A6i in combination with 5 nM selenite (right panel). Data are mean ± SEM of 3 wells of 96-well plate from 3 independent experiments. Data are normalized to vehicle control of each cell line. Cells were pre-incubated for 2 days with selenite, A6i and A6i.05 prior to RSL3 treatment under cystine containing conditions. Same control and selenite treatments were used for better comparison between control and inhibitor treatments. ** p<0.005, **** p<0.0001. Statistical analysis was performed using a two-way ANOVA with adjustment for multiple comparisons. SLC26A6 overexpressing cells treated with selenite are compared to those treated with selenite and A6i.

A6i also protected SLC26A6-overexpressing cells from increasing concentrations of selenite (Figure 3D) and improved selenite tolerance in Mel-JUSO cells expressing high endogenous SLC26A6 (Figure S7B). Furthermore, A6i selectivity was demonstrated by its failure to rescue selenite toxicity in SLC4A1-overexpressing cells, supporting its selectivity for SLC26A6 (Figure S7C). Direct measurement of intracellular selenium showed that A6i reduced selenium accumulation, whereas A6i.05 had no effect (Figure 3E). Because SLC26A6 also transports oxalate, chloride, and bicarbonate ^26,27^, we used an orthogonal assay where we measured oxalate transport. A6i reduced radiolabeled oxalate uptake in cells (Figure S7D) and inhibited oxalate–chloride exchange in SLC26A6-containing proteoliposomes in a concentration-dependent manner, with an IC_50_ in the mid-nanomolar range (Figures 3F and G). These results establish A6i as a potent and selective inhibitor of human SLC26A6-mediated anion transport.

We next examined whether SLC26A6 inhibition affects selenoprotein synthesis and ferroptosis sensitivity. A6i reduced the expression of SELENON, SELENOS, GPX1, and GPX4 in both SLC26A6-overexpressing cells and cells with high endogenous SLC26A6 expression, in the presence of selenite (Figures 3H and S7E). Consistent with reduced selenoprotein expression, A6i increased sensitivity to the GPX4 inhibitors RSL3 and FIN56, with selenium supplementation further enhancing this effect (Figures 3I and S7F). Accordingly, the inactive analog A6i.05 failed to reproduce these effects (Figures S7G and S7H). Under cysteine-replete conditions, A6i reduced GPX4 and GPX1 levels, whereas A6i.05 did not (Figure S7I). Similarly, sulfite reduced selenoprotein levels and increased sensitivity to GPX4 inhibition (Figures S7J–S7L), consistent with its ability to interfere with SLC26A6-mediated selenite uptake.

### Cryo-EM reveals the structural basis of SLC26A6 inhibition by A6i

To define the mechanism of A6i inhibition, we determined the cryo-EM structure of SLC26A6 in complex with A6i at 2.7 Å resolution (Figure S8 and Table S3). The transporter retained the characteristic SLC26 dimeric architecture, with two membrane-embedded transport units connected through cytoplasmic STAS domains ^28^ (Figure 4A). The transmembrane domains adopted an overall conformation similar to that of inhibitor-free SLC26A6 ^16^, although they were tilted approximately 5° away from the dimer axis (Figure S9A). Each transport unit comprises a rigid gate domain and a mobile core domain containing the substrate-binding site. This site includes R404, whose mutation to valine impairs transport activity ^16^ (Figures 4B and S9B). In the inward-facing conformation, this site is accessible from the cytoplasm through a water-filled cavity at the core–gate interface (Figure S9C).

**Figure 4:**
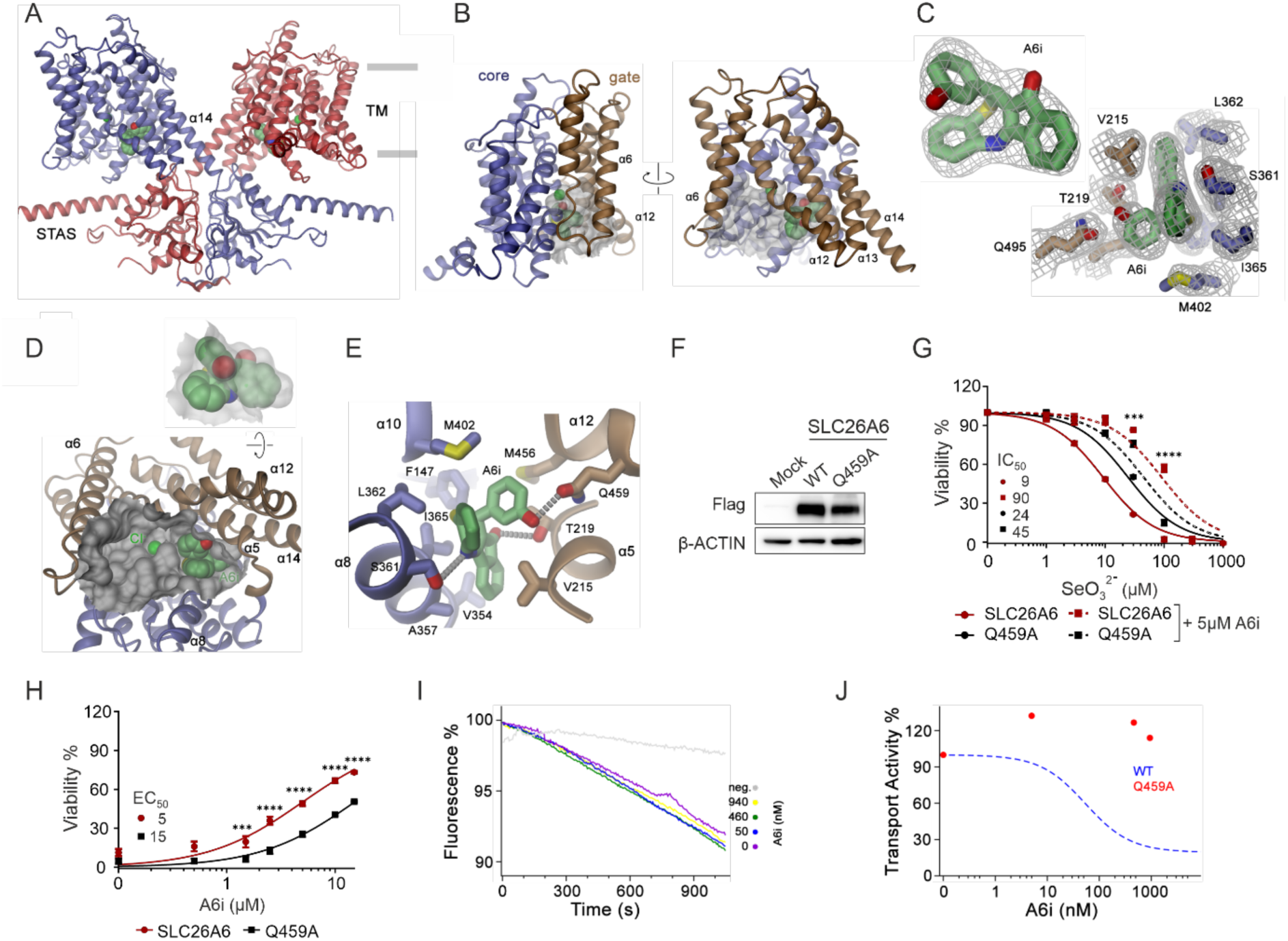
The structure of SLC26A6 bound to A6i reveals the key residues involved in the interaction. **(A)** Ribbon representation of a SLC26A6 dimer with the bound A6i occupying a site in the inward-facing cleft to the substrate binding site shown as space-filling model. The membrane boundaries are indicated with grey lines. **(B)** Ribbon representation of the transmembrane domain of a SLC26A6 subunit with the aqueous cavity leading to the anion binding site from the cytoplasm shown as surface and bound A6i as space-filling model. Core and gate domains are shown in distinctive colors. **(C)** Cryo-EM density of A6i superimposed on the model of A6i (left), and on A6i and sidechains of interacting residues (right). **(D)** Relation of bound A6i relative to the transported substrate Cl**^−^** (green). The TM is shown as ribbon in a view from the cytoplasm. The aqueous cavity leading to the ion binding site is shown as surface. Inset (top) shows view of the bound A6i with surface of the binding pocket superimposed. **(E)** Closeup of the residues surrounding the bound A6i with hydrogen bonds indicated by dashed lines. Sidechains of residues in contact with the inhibitor are shown as sticks. **(F)** Immunoblot analysis of SLC26A6 knock-out NCI-H1975 cells expressing mock, WT and Q459A mutant of SLC26A6. **(G)** Viability of SLC26A6 knock-out NCI-H1975 cells expressing mock, WT or Q459A mutants of SLC26A6, in the presence of increasing concentration of selenite, with or without 5µM of A6i. Data are mean ± SEM of 2 wells of 96-well plate from 2 independent experiments. Data are normalized to the vehicle control of each cell line. *** p<0.001, **** p<0.0001. Statistical analysis was performed using a two-way ANOVA with adjustment for multiple comparisons. SLC26A6 overexpressing cells with A6i are compared to SLC26A6-Q459A overexpressing cells with A6i. **(H)** Viability of SLC26A6 knock-out NCI-H1975 cells expressing WT or Q459A mutants of SLC26A6, upon treatment with 100µM selenite, in the presence of increasing concentration of A6i. Data are mean ± SEM of 3 wells of 96-well plate from 2 independent experiments. Data are normalized to the 500nM Lip1 treated of each cell line. *** p<0.001, **** p<0.0001. Statistical analysis was performed using a two-way ANOVA with adjustment for multiple comparisons. **(I)** Concentration-dependent inhibition of oxalate/Cl**^−^** exchange into proteoliposomes containing reconstituted SLC26A6 mutant Q459A by A6i. Transport is assayed via the parallel import of H^+^ with the pH sensitive fluorophore ACMA. Traces show means of 5 replicates from one reconstitution. **(J)** Transport activity of SLC26A6 does not decrease in presence of A6i at the assayed concentrations (red). A6i concentration-response relationships of WT (displayed in Figure 3G) are shown as dashed line for comparison.

Well-resolved density revealed A6i bound in a pocket at the edge of this cavity, near the dimer interface and distinct from the anion-binding site (Figures 4C and 4D). The inhibitor contacts residues from both the core and gate domains, including V215, T219, S361, M402, M456, and Q459, through hydrophobic and polar interactions (Figure 4E). To test the contribution of these interactions, we mutated candidate binding residues and found that substitution of Q459 with alanine reduced A6i activity in cellular selenite-toxicity assays (Figures 4F–4H). This result is consistent with the cryo-EM structure and the analysis of structure–activity relationships of related small molecules (Figure 3B), which implicated the inhibitor’s hydroxyl group in binding to Q459.

Reconstitution of purified SLC26A6-Q459A into proteoliposomes revealed reduced anion-exchange activity and no detectable inhibition by A6i across the tested concentration range (Figures 4I and 4J). Together, the structural and functional data identify the A6i-binding pocket and implicate Q459 in inhibitor recognition and SLC26A6 inhibition.

## Discussion

Our findings reported here identify SLC26A6 as a direct selenite transporter and establish a previously unrecognized route of inorganic selenium uptake in mammalian cells. By controlling selenium availability for *de novo* selenocysteine synthesis ^29,30^, SLC26A6 supports selenoprotein production, thereby promoting redox homeostasis and ferroptosis resistance. These findings link anion transport to selenium homeostasis and identify SLC26A6 as a contributor of cellular susceptibility to ferroptosis.

Our findings further suggest that extracellular cysteine availability shapes how much SLC26A6 contributes to selenium uptake and selenoprotein biosynthesis. When cysteine is abundant, SLC26A6 has a limited effect, consistent with an alternative uptake route in which cysteine reacts with selenite to form selenodicysteine, a putative substrate of system x_c_^−^. At extracellular levels found in plasma, and as cysteine becomes scarce, direct selenite uptake through SLC26A6 appears to play a larger role, helping cells to maintain selenium acquisition when extracellular thiol-dependent uptake is limited. Together, these pathways may offer complementary routes for selenium supply, with their relative contributions shaped by extracellular selenium chemistry and transporter expression. This model highlights the importance of accounting for cysteine availability when interpreting selenite supplementation experiments, since culture conditions may favor one uptake pathway over the other. However, the role of system x_c_^−^ under these conditions still requires direct functional testing, and transport of the proposed selenotrisulfide intermediate remains to be demonstrated.

Motivated by these observations, we report on the identification of A6i the first pharmacological inhibitor of human SLC26A6. We demonstrate that it reduces selenium accumulation and selenoprotein expression, thereby sensitizing cancer cell lines to ferroptosis. The pharmacological modulation of selenium uptake by SLC26A6 may have broader therapeutic implications. Given that SLC26A6 also mediates the transport of oxalate and bicarbonate, its inhibition could additionally impact conditions involving bicarbonate and oxalate homeostasis, such as kidney stone formation.

Thus, these findings establish SLC26A6-mediated selenium uptake as a potential point of intervention in selenoprotein-dependent redox regulation, while providing a framework for investigating its contribution to tumor ferroptosis resistance.

### Limitations of the Study

Our discovery of A6i represents the first human-specific inhibitor of SLC26A6. However, A6i exhibits limited activity against the murine ortholog, Slc26a6, which poses challenges for conducting preclinical studies in mouse models. To overcome this limitation, SAR studies will be necessary to optimize the compound and develop derivatives capable of inhibiting both the human and mouse isoforms.

## Acknowledgments

J.P.F.A. acknowledges the support of from Rudolf Virchow Center, University of Würzburg and additional grants from the Deutsche Forschungsgemeinschaft (DFG), FR 3746/3-1, FR 3746/5-1, FR 3746/6-1, CRC/TRR205, INST269/886-1, and TRR 387/1 (514894665); the EU-H2020 (ERC-Consolidator, DeciFERR); and the Deutsche Jose Carreras Leukämie Stiftung (DJCLS 01 R/2022). This work was supported by a grant of the Swiss National Science Foundation (10.006.508) to R.D. M.C. acknowledges support by the DFG Priority Program SPP 2306 [CO 291/9-1, #461385412; CO 291/10-1, #461507177; CO 291/9-2, CO 291/10-2, CO 291/14-1] and the European Research Council (ERC) under the European Union’s Horizon 2020 research and innovation programme (grant agreement No. GA 884754). N.L. was supported by the National Research Foundation of Korea (NRF) grant funded by the Korea government (MSIT)(RS-2024-00338061).

Generative artificial intelligence assisted with language editing. The authors are responsible for the accuracy and integrity of the final manuscript.

## Author Contributions

S.A. and G.F. performed most of the experiments and analyzed data. M.S., and J.G. contributed to the development of the methodology and performed experiments. Z.C., and A.F.D.S performed the genetic screens and analyzed related data. P.K. purified and reconstituted SLC26A6 constructs, performed *in vitro* transport experiments, collected cryo-EM data, and determined the structure of the SLC26A6-A6i inhibitor complex. D.T. supported protein production and cryo-EM data collection. R.D. supervised structural and *in vitro* experiments and contributed to data analysis and manuscript preparation. M.Sch., and A.P.K. performed TXRF. M.F. provided funding and guidance. A.C.N., and F.K. contributed to C14 oxalate measurements in combination with A6i. N.L. contributed to selenium measurement. K.K., M.E., H.N., N.G. and H.A. performed the *in vivo* experiments with the neuroblastoma implants. M.S., S.A., C.D., B.P., E.M., and M.C. performed small molecule library screen to identify the A6i. E.G., and G.S-F provided SLC-focused KO library and constructs of SLC26A family. J.P.F.A. conceived the study, designed the experimental plan, and supervised the work. G.F, S.A., and J.P.F.A. wrote the first version of the manuscript. The final version received input from all authors. All authors contributed to data discussion, interpretation, revision and approved the final version of the manuscript.

## Competing interests

J.P.F.A., S.A., M.C., B.P., and E.M. hold patent application for A6i compound described in this study. M.C. is a co-founder and shareholder of ROSCUE Therapeutics GmbH.

---

**Figure S1:**
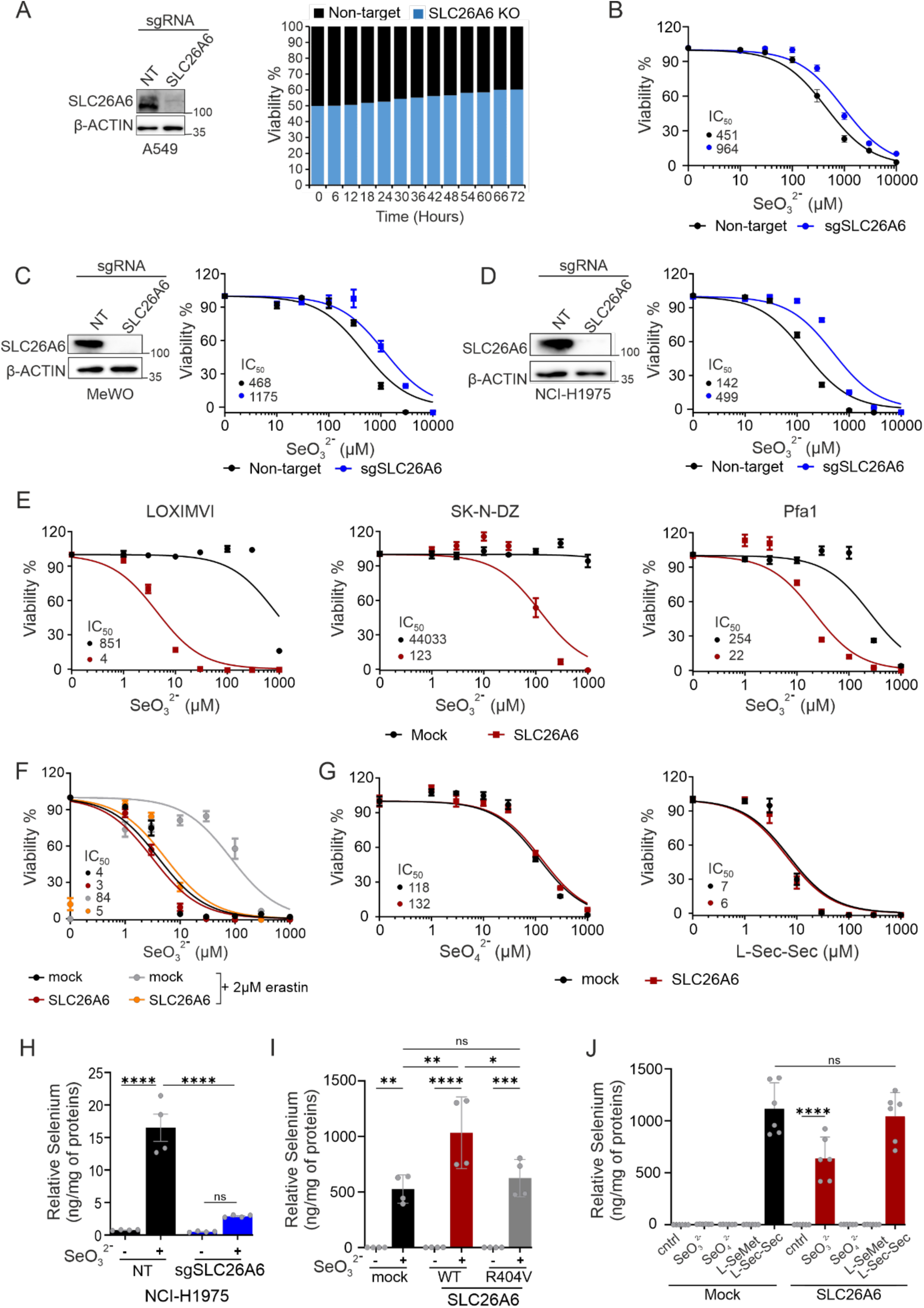
Knocking out SLC26A6 leads to resistance to sodium selenite treatment. **(A)** Immunoblot analysis and competition of A549 cells transduced with non-target or SLC26A6 sgRNA in the presence of 30 µM of sodium selenite in normal medium. Data are mean ± SEM of 3 wells of 96-well plate from 2 independent experiments. Data are normalized to time zero. **(B)** Viability of A549 cells transduced with non-target or SLC26A6 sgRNA in the presence of increasing concentration of sodium selenite in the cystine free medium. Data are the mean ± SEM of 3 wells of 96-well plate from 3 independent experiments. Data are normalized to vehicle control of each cell line. **(C)** Immunoblot and viability of MeWo cells transduced with non-target or SLC26A6 sgRNA in the presence of increasing concentration of sodium selenite in the cystine free medium. Data are the mean ± SEM of 3 wells of 96-well plate from 2 independent experiments. Data are normalized to vehicle control of each cell line. **(D)** Immunoblot and viability of NCI-H1975 cells transduced with non-target or SLC26A6 sgRNA in the presence of increasing concentration of sodium selenite in the cystine free medium. Data are the mean ± SEM of 3 wells of 96-well plate from 2 independent experiments. Data are normalized to vehicle control of each cell line. **(E)** Viability of LOXIMVI, SK-N-DZ and Pfa1 cells transduced with mock or SLC26A6 overexpression vector in the presence of increasing concentrations of sodium selenite. Data are the mean ± SEM of 3 wells of 96-well plate from 1 (LOXIMVI) or 3 independent experiments (SK-N-DZ and Pfa1). Data are normalized to vehicle control of each cell line. **(F)** Viability of HT1080 cells transduced with mock or SLC26A6 overexpression vector in the cystine-containing medium and in the presence of increasing concentrations of sodium selenite, with or without the SLC7A11 inhibitor, erastin. Data are the mean ± SEM of 2 wells of 96-well plate from 3 independent experiments. Data are normalized to vehicle control of each cell line. **(G)** Viability of HT1080 cells transduced with mock or SLC26A6 overexpression vector, in the presence of increasing concentration of sodium selenate or selenocysteine (L-sec-sec). Data are the mean ± SEM of 3 wells of 96-well plate from 2 independent experiments. Data are normalized to vehicle control of each cell line. **(H)** Total selenium content of NCI-H1975 cells transduced with non-target or SLC26A6 sgRNA with or without 50 μM of selenite treatment for 6 hours. **** p<0.0001. Statistical analysis was performed using a two-way ANOVA with adjustment for multiple comparisons. **(I)** Total selenium content of HT1080 cells transduced with mock, WT or R404V mutant of SLC26A6 in cystine-containing medium with or without 50µM selenite treatment. * p<0.05, ** p<0.005, *** p<0.0005, **** p<0.0001. Statistical analysis was performed using a two-way ANOVA with adjustment for multiple comparison. **(J)** Total selenium content of SLC26A6 overexpressing HT1080 cells treated with 50µM selenite, selenate, selenomethionine, or 25µM selenocystine in cysteine-free medium. **** p<0.0001. Statistical analysis was performed using a two-way ANOVA with adjustment for multiple comparison.

**Figure S2:**
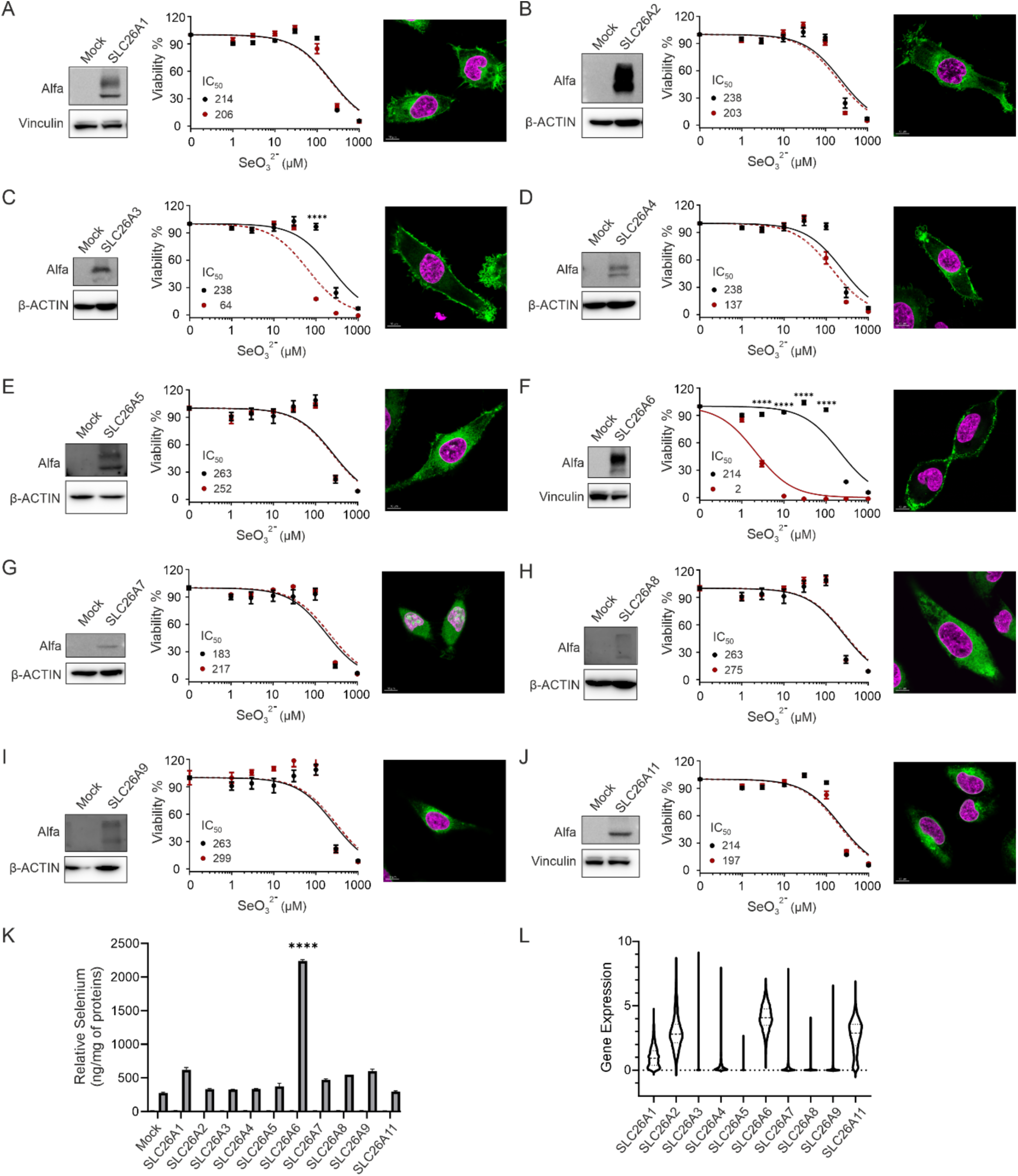
SLC26A6 overexpression promotes sensitivity to sodium selenite more efficiently compared to other SLC26A family members. Immunoblot analysis, viability, and confocal microscopy images of HT1080 cells transduced with mock or overexpression vectors for the indicated SLC26A family members. The expression of each construct was confirmed by blotting for the N-terminally conjugated Alfa-tag for SLC26A1 **(A)**, SLC26A2 **(B)**, SLC26A3 **(C)**, SLC26A4 **(D)**, SLC26A5 **(E)**, SLC26A6 **(F)**, SLC26A7 **(G)**, SLC26A8 **(H)**, SLC26A9 **(I)**, SLC26A11 **(J)**. The viability is assessed in the presence of increasing concentrations of sodium selenite. Subcellular localization of each transporter is shown by Alfa-tag staining (green). DNA is counterstained with DAPI (magenta). Data are the mean ± SEM of 3 wells of 96-well plate from 3 independent experiments. For SLC26A5, SLC26A7, SLC26A8 and SLC26A9, data are the mean ± SD of 3 wells of 96 well plate from 1 experiment. **** p<0.0001. Statistical analysis was performed using a two-way ANOVA with adjustment for multiple comparison. **(K)** Total selenium content of HT1080 cells transduced with mock or the indicated members of SLC26A family after treatment with 50 μM of selenite for 6 hours. Total selenium was measured using TXRF method. 1mg/L Yttrium was used as standard. Measurements are normalized to protein levels. **** p<0.0001. Statistical analysis was performed using a two-way ANOVA with adjustment for multiple comparisons. **(L)** Gene expression profile of the indicated members of the SLC26A family across different tissues in human samples. mRNA expression values are derived from DepMap (https://depmap.org).

**Figure S3:**
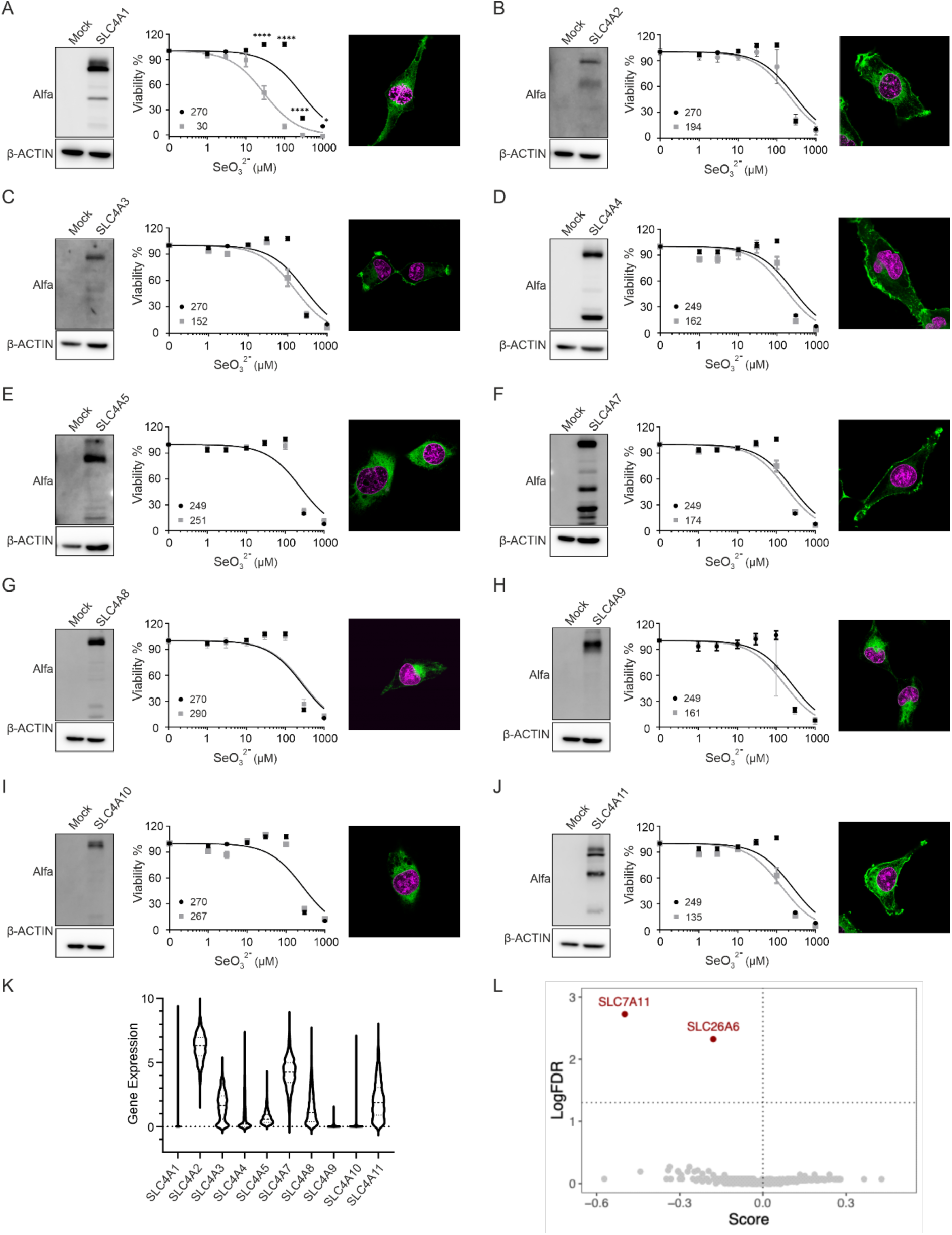
Impact of the SLC4A family and other transporters on selenite-induced toxicity. Immunoblot analysis, viability and confocal microscopy images of HT1080 cells transduced with mock or overexpression vectors for the indicated SLC4A family members. The expression of each construct was confirmed by blotting for the N-terminally conjugated Alfa-tag for SLC4A1 **(A)**, SLC4A2 **(B)**, SLC4A3 **(C)**, SLC4A4 **(D)**, SLC4A5 **(E)**, SLC4A7 **(F)**, SLC4A8 **(G)**, SLC4A9 **(H)**, SLC4A10 **(I)**, SLC4A11 **(J)**. The viability is assessed in the presence of increasing concentrations of sodium selenite. Subcellular localization of each transporter is shown by Alfa-tag staining (green). DNA is counterstained with DAPI (magenta). Data are the mean ± SEM of 3 wells of 96-well plate from 2 independent experiments. * p<0.05, **** p<0.0001. Statistical analysis was performed using a two-way ANOVA with adjustment for multiple comparison. **(K)** Gene expression profile of the indicated members of the SLC4A family across different human cell lines. mRNA expression values are derived from DepMap (https://depmap.org). **(L)** LogFDR results for CRISPR activation screen in HT1080-vp64 dCas9-MPH2 v2 cells upon supplementation with 50µM of selenite, showing SLC7A11 and SLC26A6 as top hits.

**Figure S4:**
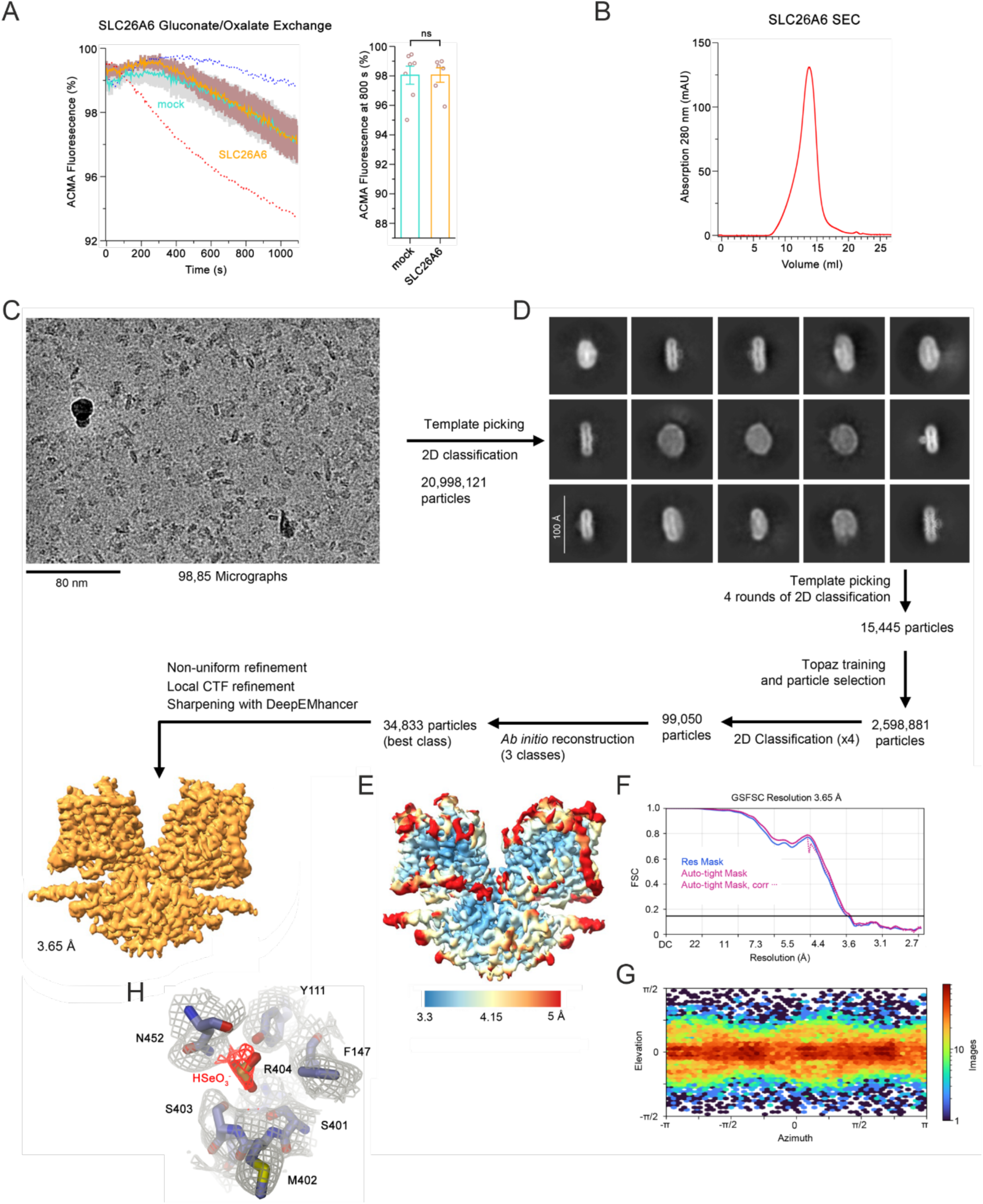
SLC26A6 selenite interaction and transport. **(A)** Equivalent experiments as shown in A probing the exchange of oxalate with gluconate, which is not transported by SLC26A6 as a control. Proteoliposomes containing reconstituted SLC26A6 are shown in comparison to mock liposomes. Left, time dependence of ion transport. Traces show means of 6 replicates from 2 independent reconstitutions for SLC26A6 containing proteoliposomes and 7 replicates from 2 independent reconstitutions for mock. Average traces of oxalate/selenite exchange (A) are shown as dashed lines as reference. Right, comparison of the relative fluorescence decrease after 800 Sec. Individual values are indicated. The difference between SLC26A6 and mock liposome is not statistically significant. Error bars in A and B represent SEM. Statistical analysis was performed using a Welch’s ANOVA. **(B)** Size exclusion chromatography trace of SLC26A6 purified in 150 mM gluconate in the absence of transported anion on a Superose 6 column. Peak fractions containing high concentrations of the transporter were concentrated, supplemented with 50mM selenite and used for cryo-EM data collection. **(C)** Representative micrograph (of a total of 98,850) of the SLC26A6-selenite dataset. **(D)** 2D class averages of the SLC26A6-selenite complex and data-processing workflow performed in Cryo-SPARC yielding a map at 3.65 Å resolution. **(E)** Final 3D reconstruction colored according to the local resolution. **(F)** FSC plot of cryo-EM density maps of the SLC26A6/selenite complex used for the estimation of its resolution. The dotted line indicates the resolution at which the FSC drops below the 0.143 threshold. **(G)** Heatmap displaying the angular distribution of particle orientations. **(H)** Section of the map around the substrate binding site in the core domain of SLC26A6. The final cryo-EM map (contoured at 3σ) is shown superimposed on displayed residues of the refined structure. Density surrounding a selenite ion is colored in red.

**Figure S5:**
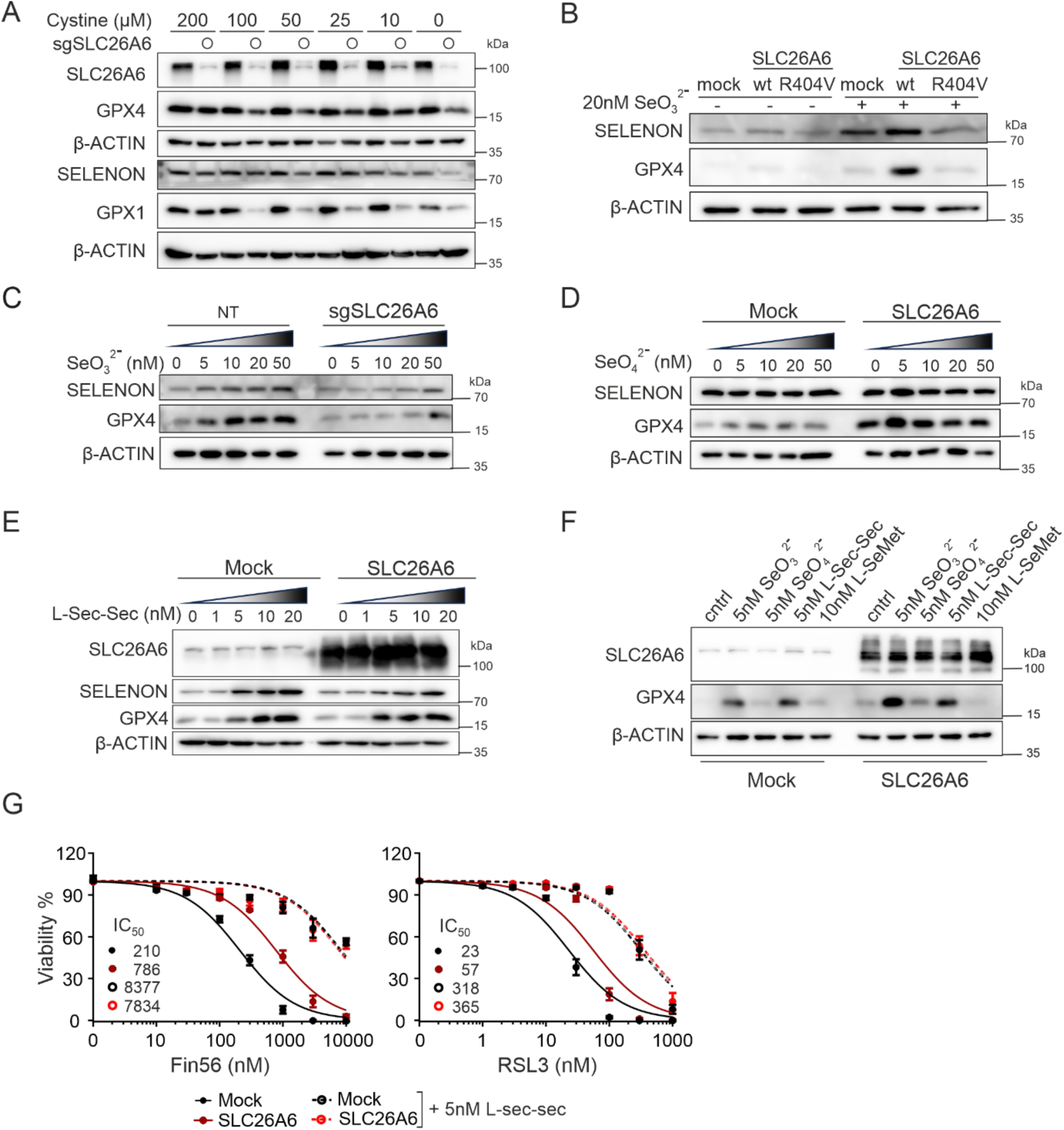
System xc− contributes to selenite uptake, while SLC26A6 transports selenium exclusively in the form of selenite. **(A)** Immunoblot analysis of the selenoproteins GPX4, GPX1, and SELENON in response to 20 nM of selenite, 500nM Lip1, and indicated concentrations of cystine in mock or SLC26A6 overexpressing NCIH1975 cells. **(B)** Immunoblot analysis of SELENON and GPX4 in response to 20 nM selenite in HT1080 cells transduced with mock, WT or R404V mutant of SLC26A6. **(C)** Immunoblot analysis of SELENON and GPX4 in response to increasing nanomolar concentrations of selenite in Mel-JUSO cells transduced with non-target or sgRNA targeting SLC26A6 in cystine-free condition. **(D-E)** Immunoblot analysis of the selenoproteins SELENON and GPX4 in response to increasing nanomolar concentrations of selenate (**D**) or selenocysteine (**E**) in mock or SLC26A6 overexpressing HT1080 cells. **(F)** Immunoblot analysis of the selenoproteins SELENON and GPX4 in response to low concentrations of selenium sources in mock or SLC26A6 overexpressing HT1080 cells. **(G)** Viability of HT1080 cells transduced with mock or SLC26A6 overexpression vector in the presence of increasing concentration of GPX4 inhibitor Fin56 and RSL3, in combination with selenocystine. Data represents mean ± SEM from 3 independent experiments. Cells were pre-incubated for 2 days with selenocystine before treatment with GPX4 inhibitor in normal medium.

**Figure S6:**
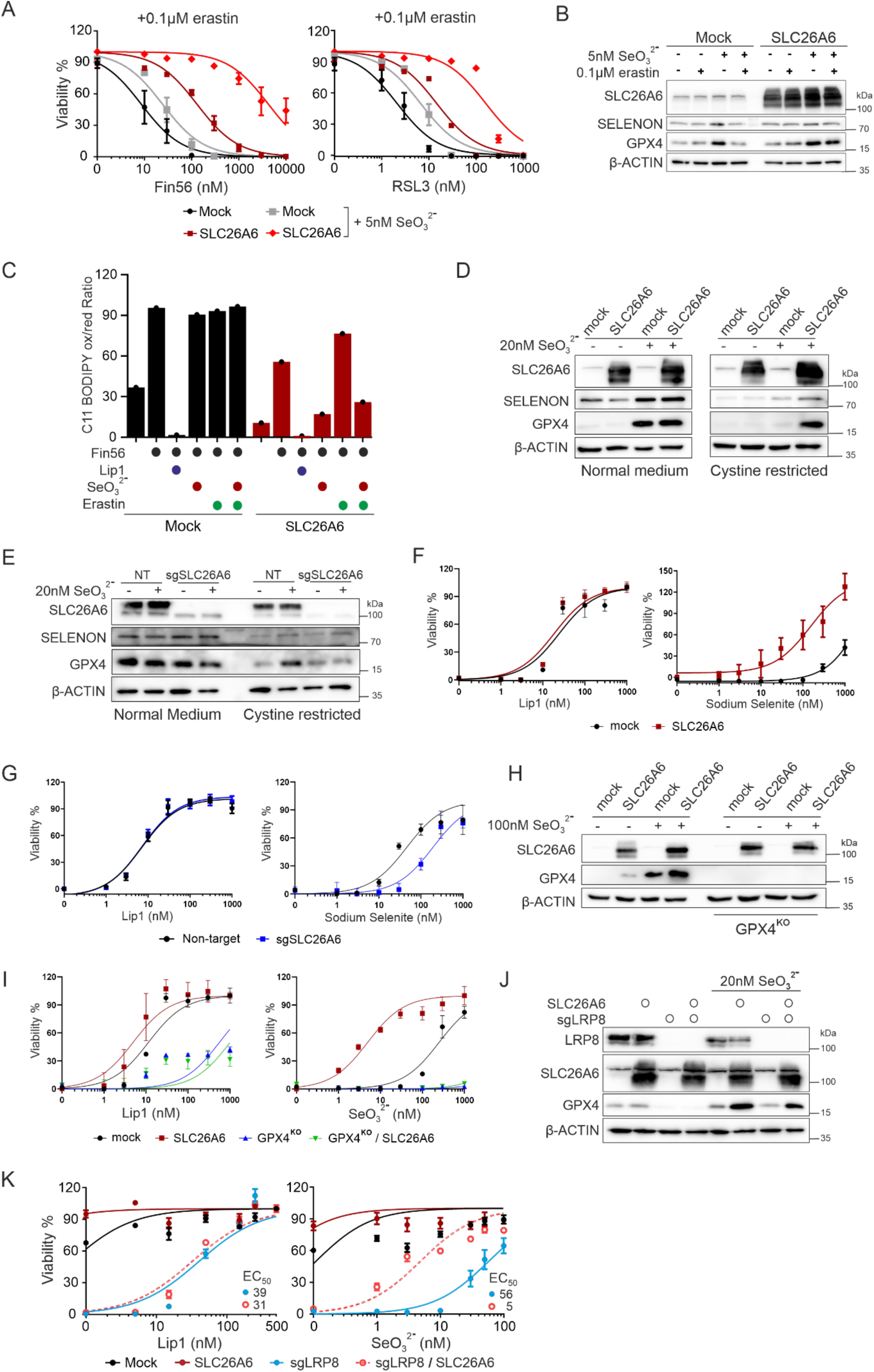
SLC26A6 overexpression protects the cells against ferroptosis induction. **(A)** Viability of HT1080 cells transduced with mock or SLC26A6 overexpression vector in the presence of increasing concentration of GPX4 inhibitor Fin56 and RSL3, in combination with selenite and erastin treatment. Data are mean ± SEM of 2 wells of 96-well plate from 3 experiments. Data are normalized to vehicle control of each cell line. Cells were pre-incubated for 2 days with selenite and erastin before treatment with GPX4 inhibitor in normal medium. **(B)** Immunoblot analysis of the selenoproteins SELENON and GPX4 in response to 5 nM selenite treatment in mock and SLC26A6 overexpressing cells with or without erastin in cystine containing medium. **(C)** Analysis of C11-BODIPY 581/591 oxidation in HT1080 cells transduced with mock, WT or R404V mutant of SLC26A6 upon treatment with Fin56, low concentration of selenite (5nM) and 0.1 µM erastin. Data represents mean ± SEM from one experiment. **(D)** Immunoblot analysis of the selenoproteins SELENON and GPX4 in response to 20 nM selenite in mock or SLC26A6 overexpressing HT1080 cells in normal and cystine free medium. **(E)** Immunoblot analysis of SELENON and GPX4 in response to 20 nM selenite in Mel-JUSO cells transduced with non-target or sgRNA targeting SLC26A6 in normal and cystine free medium. **(F)** Viability of HT1080 cells, transduced with mock or SLC26A6 overexpressing vector in the presence of increasing concentrations of liproxstatin1 (left) or sodium selenite (right) in the cysteine-free medium. Data are the mean ± SEM of 3 wells of 96-well plate from 2 independent experiments. Data are normalized to lip1 treated mock and SLC26A6 overexpressing cells. **(G)** Viability of Mel-JUSO cells, transduced with non-target or sgRNA targeting SLC26A6 in the presence of increasing concentrations of liproxstatin1 (left) or sodium selenite (right) in the cysteine-free medium. Data are the mean ± SEM of 3 wells of 96-well plate from 2 independent experiments. Data are normalized to lip1 treated mock and SLC26A6 knock-out cells. **(H)** Immunoblot analysis of WT or GPX4 knockout HT1080 cells transduced with mock or SLC26A6 overexpression vector with or without treatment with 100 nM sodium selenite. **(I)** Viability of cells from panel F in the presence of increasing concentrations of the ferroptosis inhibitor liproxstatin1 (Lip1, left) or sodium selenite (right). Data are mean ± SEM of 3 wells of 96-well plate from 1 experiment. Data are normalized to lip1 treated mock and SLC26A6 overexpressed cells. **(J)** Immunoblot analysis of SK-N-DZ cells transduced with SLC26A6 overexpression vector and/or sgRNA targeting LRP8, with or without treatment with 20 nM of sodium selenite. **(K)** Viability of SK-N-DZ cells transduced with SLC26A6 overexpression vector and/or sgRNA targeting LRP8, in the presence of increasing concentration of Lip-1 (left) or sodium selenite (right) in dialyzed serum. Data are mean ± SEM of 3 wells of 96-well plate from 2 independent experiments. Data are normalized to 500nM lip1 treatment of each cell line.

**Figure S7:**
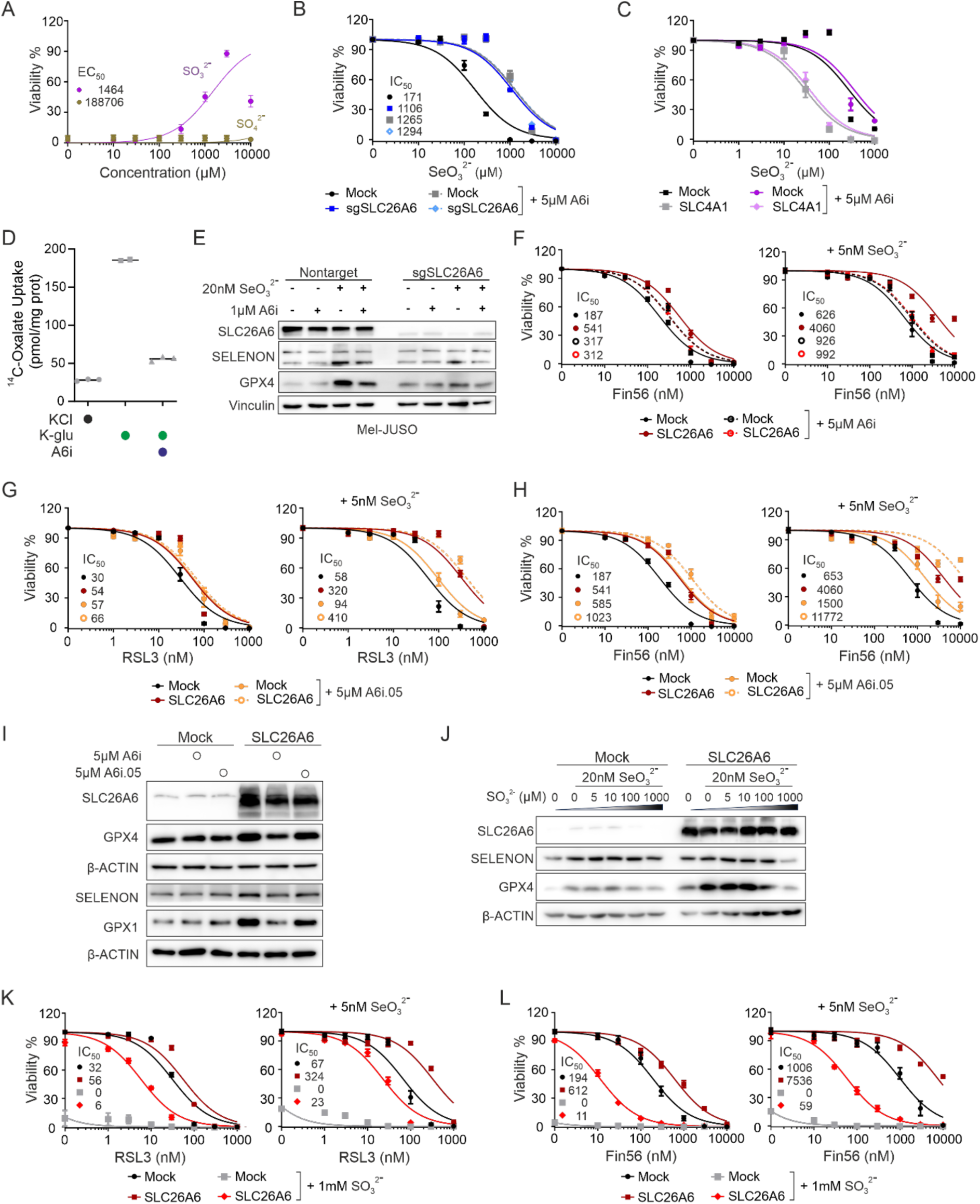
A6i and sulfite treatment reverses selenite-induced ferroptosis resistance. **(A)** Viability of SLC26A6 overexpressing cells treated with 50 µM of sodium selenite together with the increasing concentration of sulfite and sulfate. Data are the mean ± SEM of 3 wells of 96-well plate from 3 independent experiments. Data are normalized to vehicle control of each cell line. **(B)** Viability of mock and SLC26A6 knockout Mel-JUSO cells, in the presence of increasing concentration of selenite, with or without 5µM of A6i. Data are mean ± SEM of 3 wells of 96-well plate from 2 independent experiments. Data are normalized to the vehicle control of each cell line. **(C)** Viability of mock and SLC4A1 overexpressing cells, in the presence of increasing concentration of selenite, with or without 5µM of A6i. Data are mean ± SEM of 3 wells of 96-well plate from 3 independent experiments. Data are normalized to the vehicle control of each cell line. **(D)** ^14^C labelled oxalate uptake measurements, performed in the presence of KCl, k-glutarate with or without A6i treatment. **(E)** Immunoblot analyses of the selenoproteins SELENON and GPX4 in response to 1µM A6i in combination with 20 nM selenite in MEL-JUSO cells transduced with nontarget or sgRNA targeting SLC26A6. **(F)** Viability of mock and SLC26A6 overexpressing HT1080 cells in the presence of increasing concentration of Fin56 with or without A6i in combination with 5 nM selenite (right panel). Data are mean ± SEM of 2 wells of 96-well plate from 3 independent experiments. Data are normalized to vehicle control of each cell line. Cells were pre-incubated for 2 days with selenite and A6i prior to Fin56 treatment under cystine containing conditions. Same control and selenite treatments were used for better comparison between control and inhibitor treatments in (**F**) and (**H**). **(G-H)** Viability of mock and SLC26A6 overexpressing HT1080 cells in the presence of increasing concentration of RSL3 **(G)** or Fin56 **(H)** with or without A6i.05, in combination with 5 nM selenite (right panels). Data are mean ± SEM of 3 wells of 96-well plate from 3 independent experiments. Data are normalized to vehicle control of each cell line. Cells were pre-incubated for 2 days with selenite and A6i.05 prior to Fin56 or RSL3 treatment under cystine containing conditions. Same control and selenite treatments were used for better comparison between control and inhibitor treatments in (**F**) and (**H**). **(I)** Immunoblot analysis of the selenoproteins GPX4, GPX1 and SELENON in response to 5µM A6i or A6i.05 in mock or SLC26A6 overexpressing HT1080 cells in cysteine containing conditions. **(J)** Immunoblot analysis of the selenoproteins SELENON and GPX4 in response to increasing concentrations of sulfite in combination with 20nM selenite in mock or SLC26A6 overexpressing HT1080 cells. **(K-L)** Viability of mock or SLC26A6 overexpressing HT1080 cells in the presence of increasing concentration of the GPX4 inhibitor RSL3 **(K)** or Fin56 **(L)** together with or without sodium selenite and 1 mM sulfite. Data are mean ± SD of 3 wells of 96-well plate from 1 experiment. Data are normalized to vehicle control of each untreated cell line. Cells were pre-incubated for 2 days with selenite and sulfite prior to RSL3 or Fin56 treatment under cystine containing conditions.

**Figure S8:**
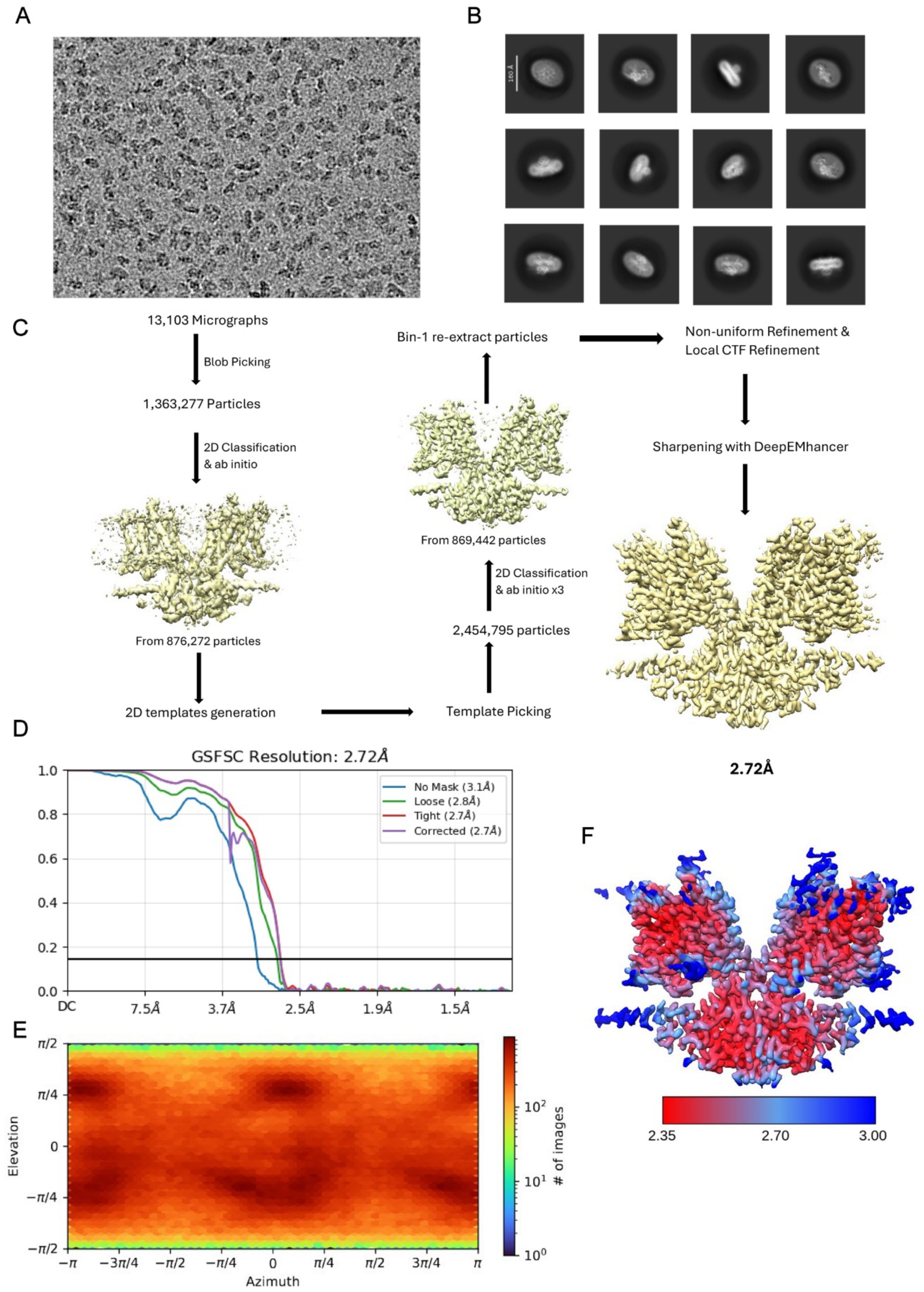
Cryo-EM reconstruction of the SLC26A6/A6i complex. **(A)** Representative micrograph (of a total of 13,103) of the SLC26A6-A6i dataset. **(B)** 2D class averages of the SLC26A6-A6i complex. **(C)** Data-processing workflow performed in Cryo-SPARC yielding a map at 2.72 Å resolution. After motion correction and CTF estimation, blob picking yielded 1,363,277 particles. Particles were subjected to 2D classification and *ab initio* reconstruction, providing a subset of 876,272 particles. 2D templates were generated and used for template-based picking, followed by 3 rounds of 2D classification and *ab initio* reconstruction with C2 symmetry applied, yielding a density map from 869,442 particles. These were re-extracted (after unbinning) and the new map was improved by non-uniform refinement, local CTF refinement and DeepEMhancer sharpening, reaching a resolution of 2.72 Å. **(D)** FSC plot of cryo-EM density maps of the SLC26A6-A6i complex used for the estimation of its resolution. The dotted line indicates the resolution at which the FSC drops below the 0.143 threshold. **(E)** Heatmap displaying the angular distribution of particle orientations. **(F)** Final 3D reconstruction colored according to the local resolution.

**Figure S9:**
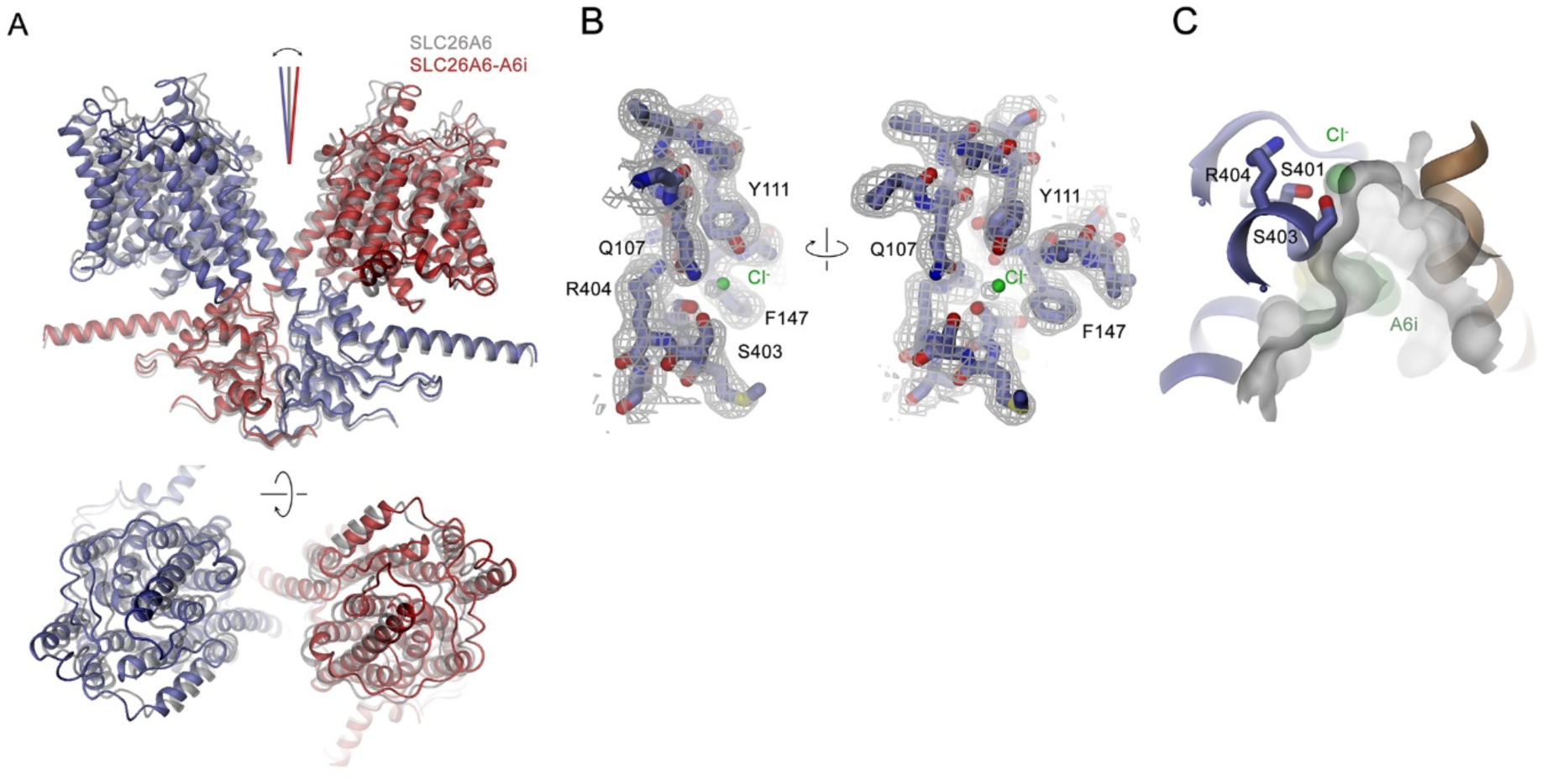
Structural features of the SLC26A6-A6i complex. **(A)** Superposition of the structure of the SLC26A6-A6i complex (red, blue) on the SLC26A6 dimer (grey) shows an outward tilt of the TM units by 5° from the dimer axis. Top, view from within the membrane with the extracellular side on top. Bottom, view from the extracellular side. **(B)** Cryo-EM density of the ion binding site located in the core domain of the TM subunits superimposed on the refined model. A chloride ion is shown as green sphere. **(C)** Section of the molecular surface of the aqueous cavity leading from the cytoplasm to the ion binding site of the TM unit superimposed on a ribbon of the protein. Selected residues coordinating the ion are shown as sticks.

**Table S1.** Structure-activity relationship of A6i analogs.

| **Compound** | **R1** | **R2** | **IC<sub>50</sub>** |
|---|---|---|---|
| **A6i** |  |  | **2.8** |
| **A6i.01** |  |  | **15** |
| **A6i.02** |  |  | **>15** |
| **A6i.03** |  |  | **>15** |
| **A6i.04** |  |  | **>15** |
| **A6i.05** |  |  | **>15** |
| **A6i.06** |  |  | **>15** |
| **A6i.07** |  |  | **>15** |
| **A6i.08** |  |  | **>15** |
| **A6i.09** |  |  | **>15** |
| **A6i.10** |  |  | **>15** |
| **A6i.11** |  |  | **>15** |
| **A6i.12** |  |  | **>15** |
| **A6i.13** |  |  | **>15** |

## Materials & Method

## STAR Methods

### Key Resources

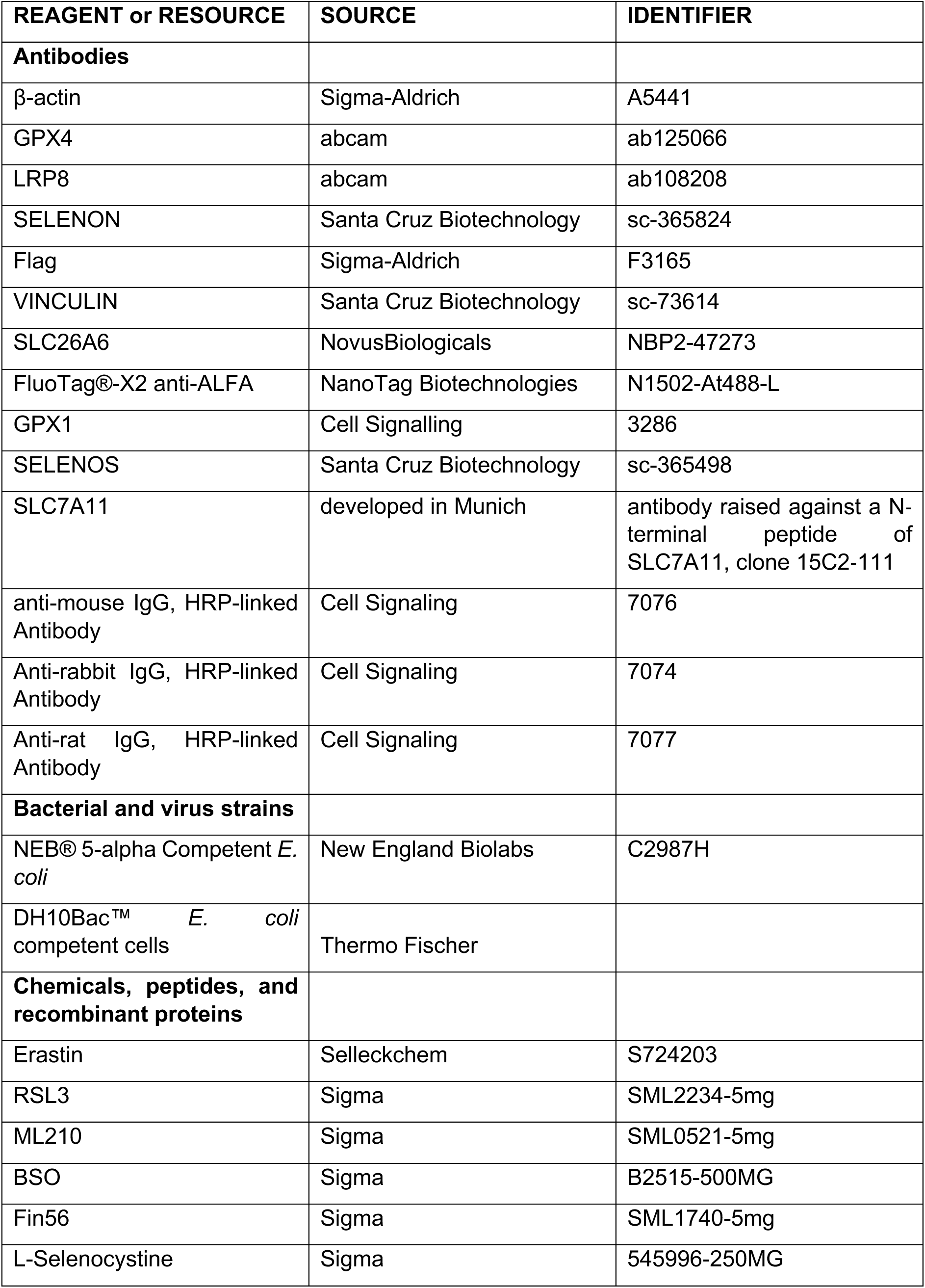
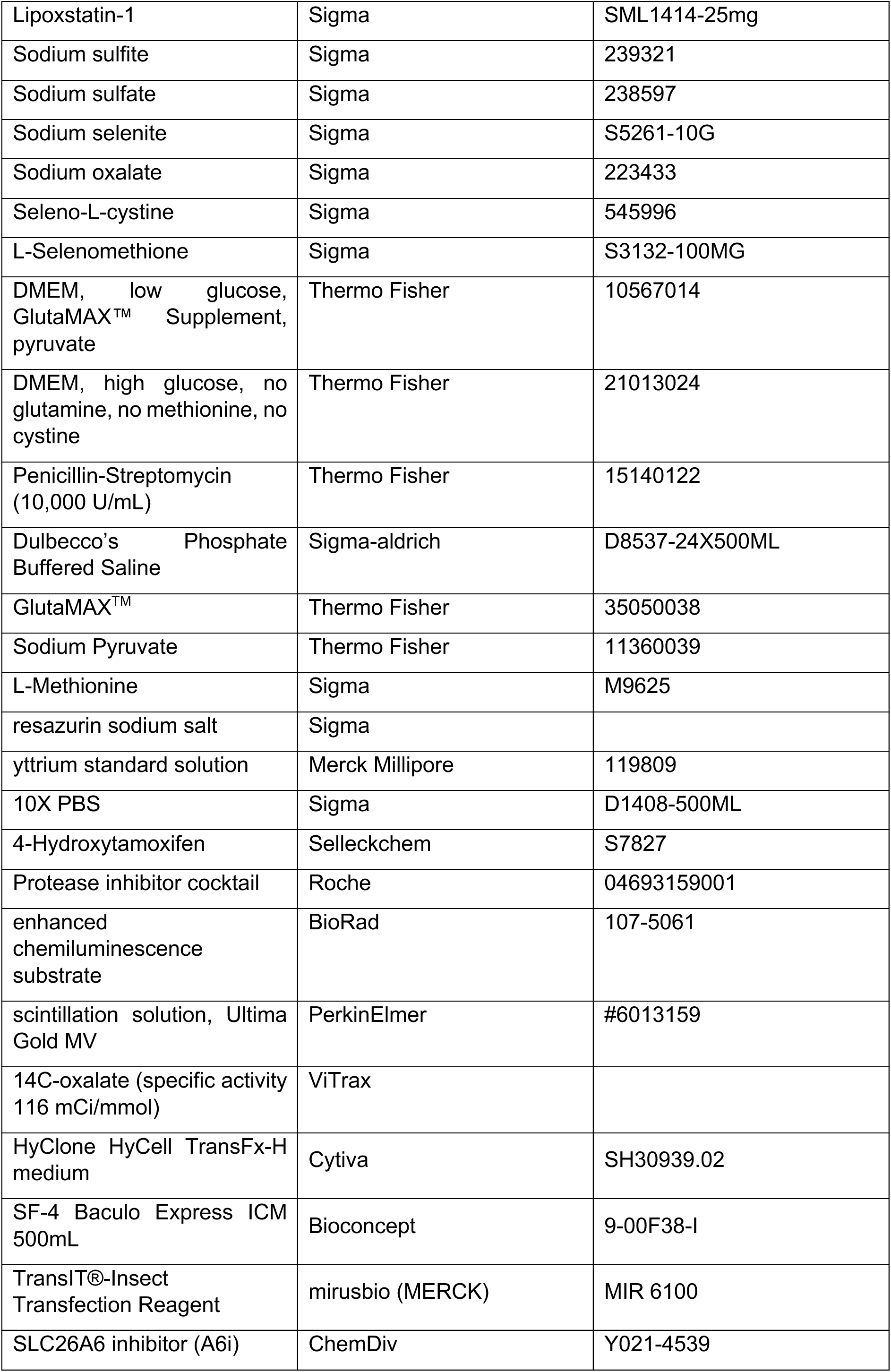
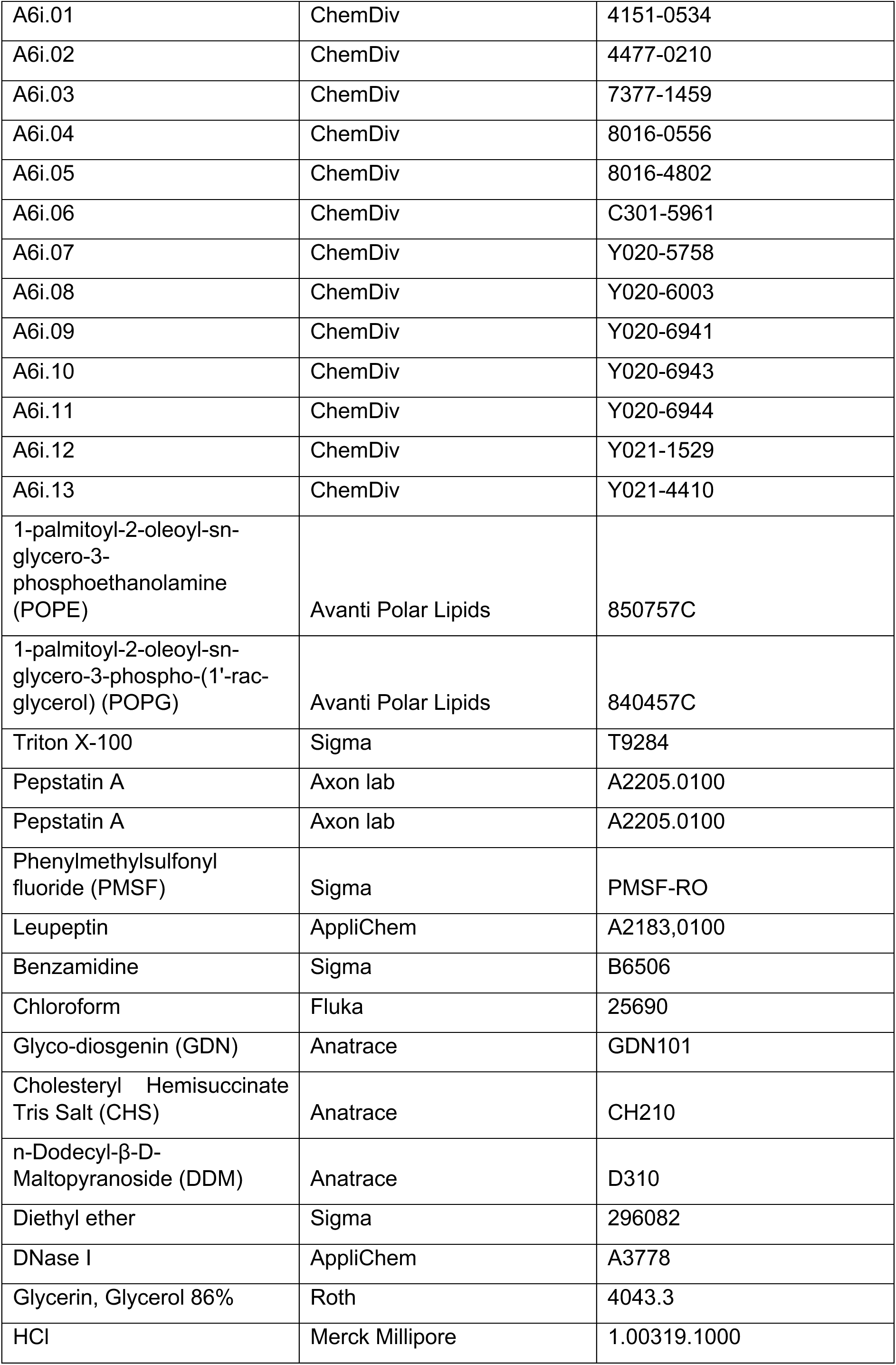
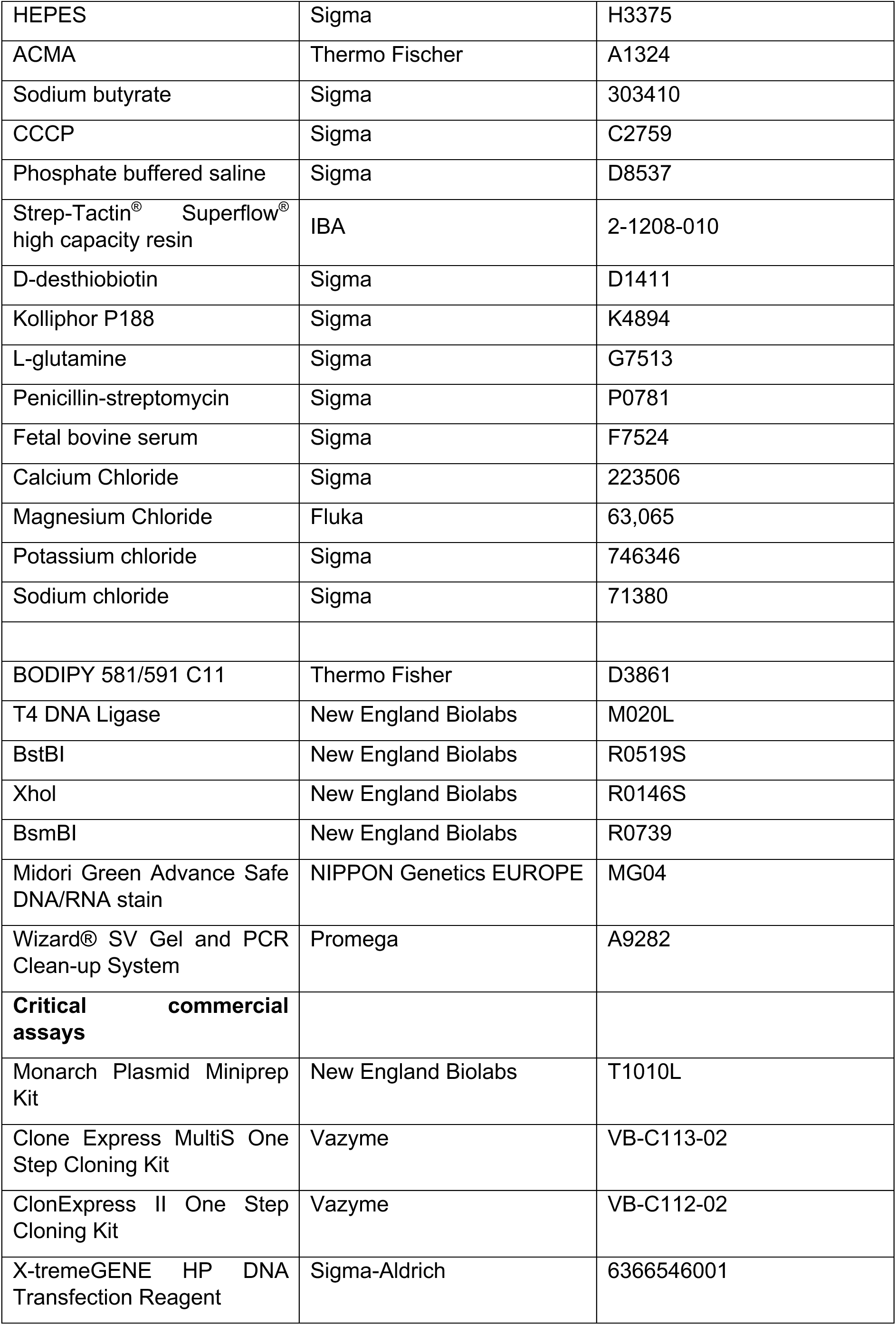
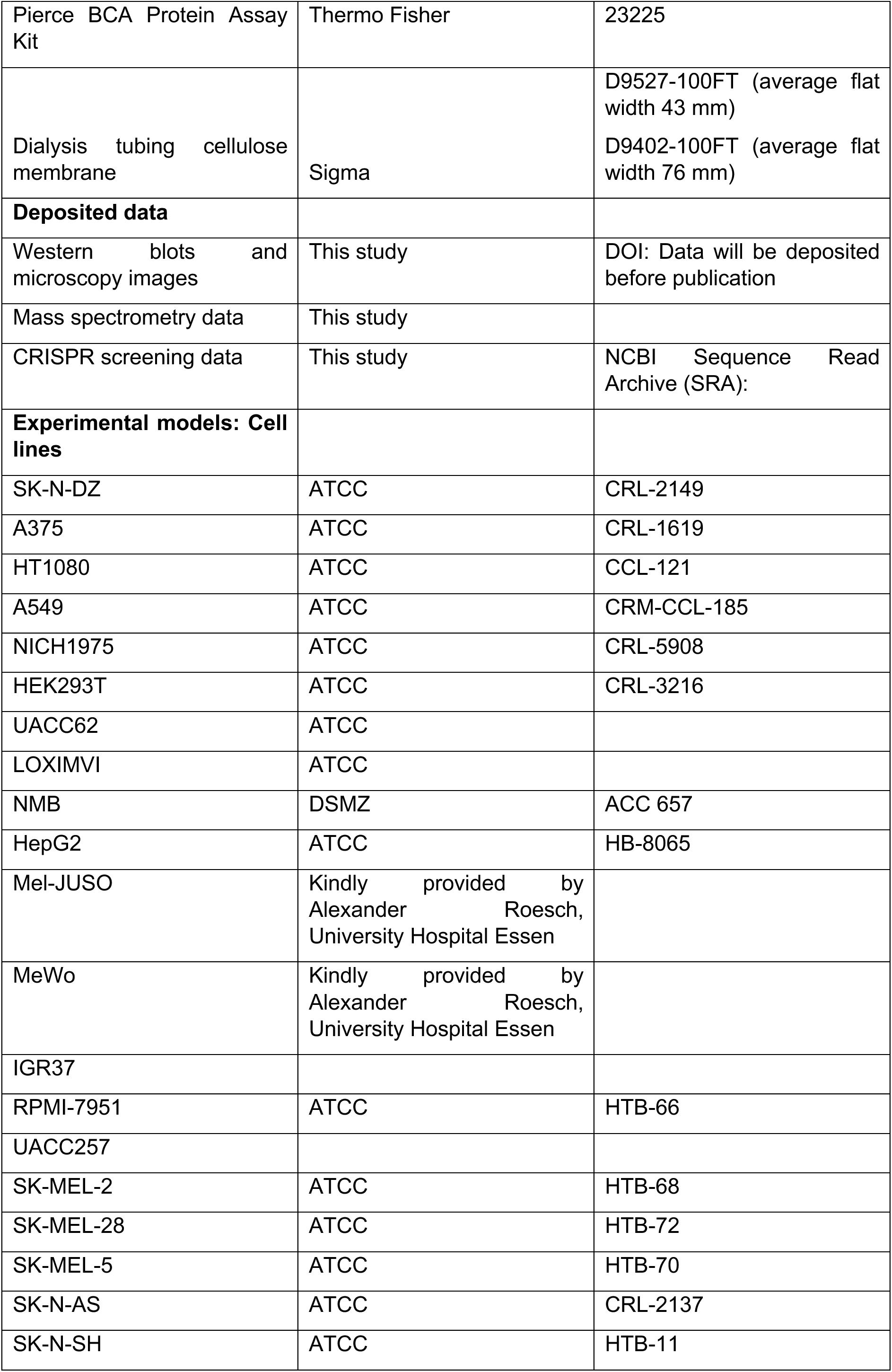
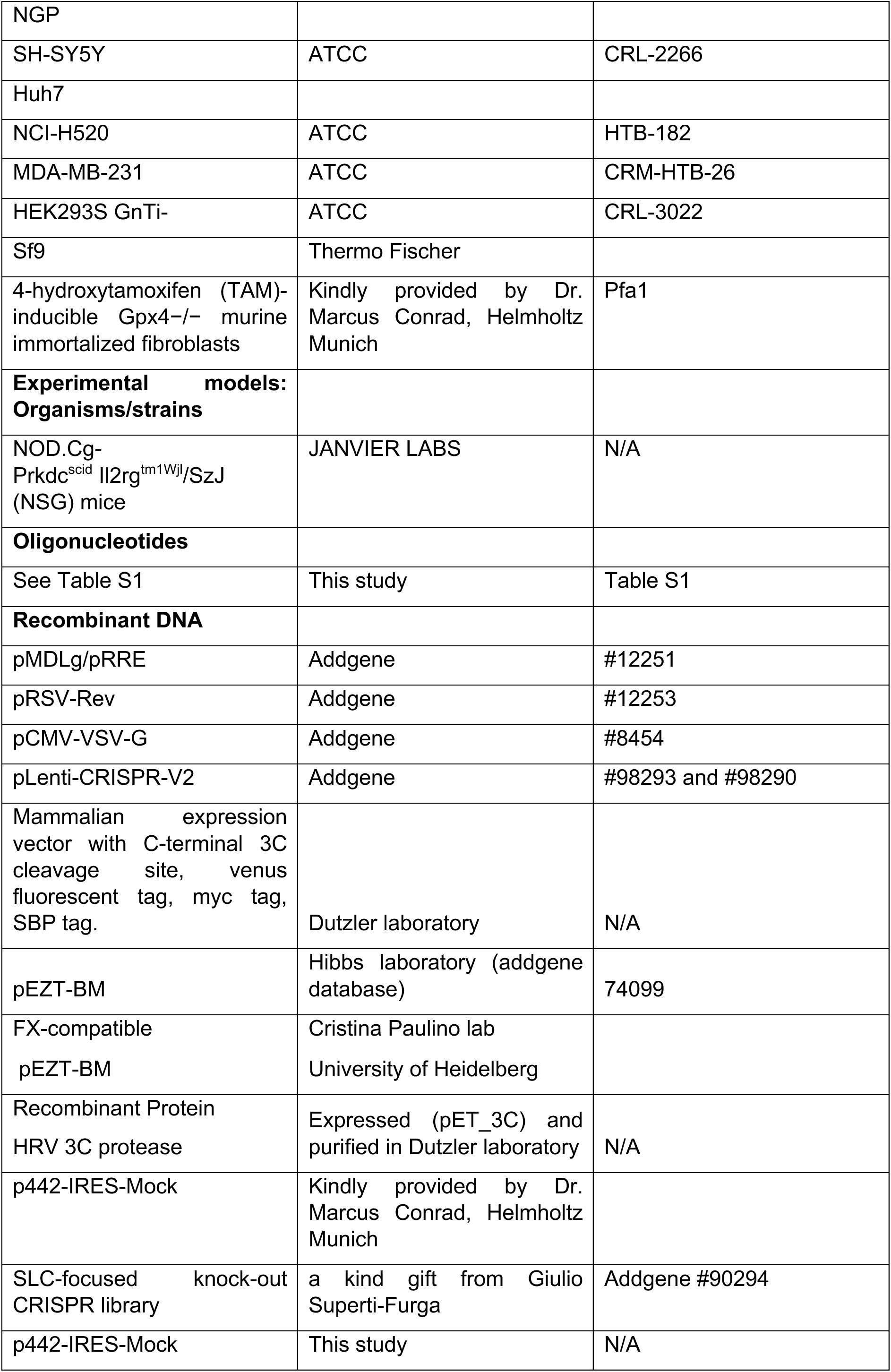
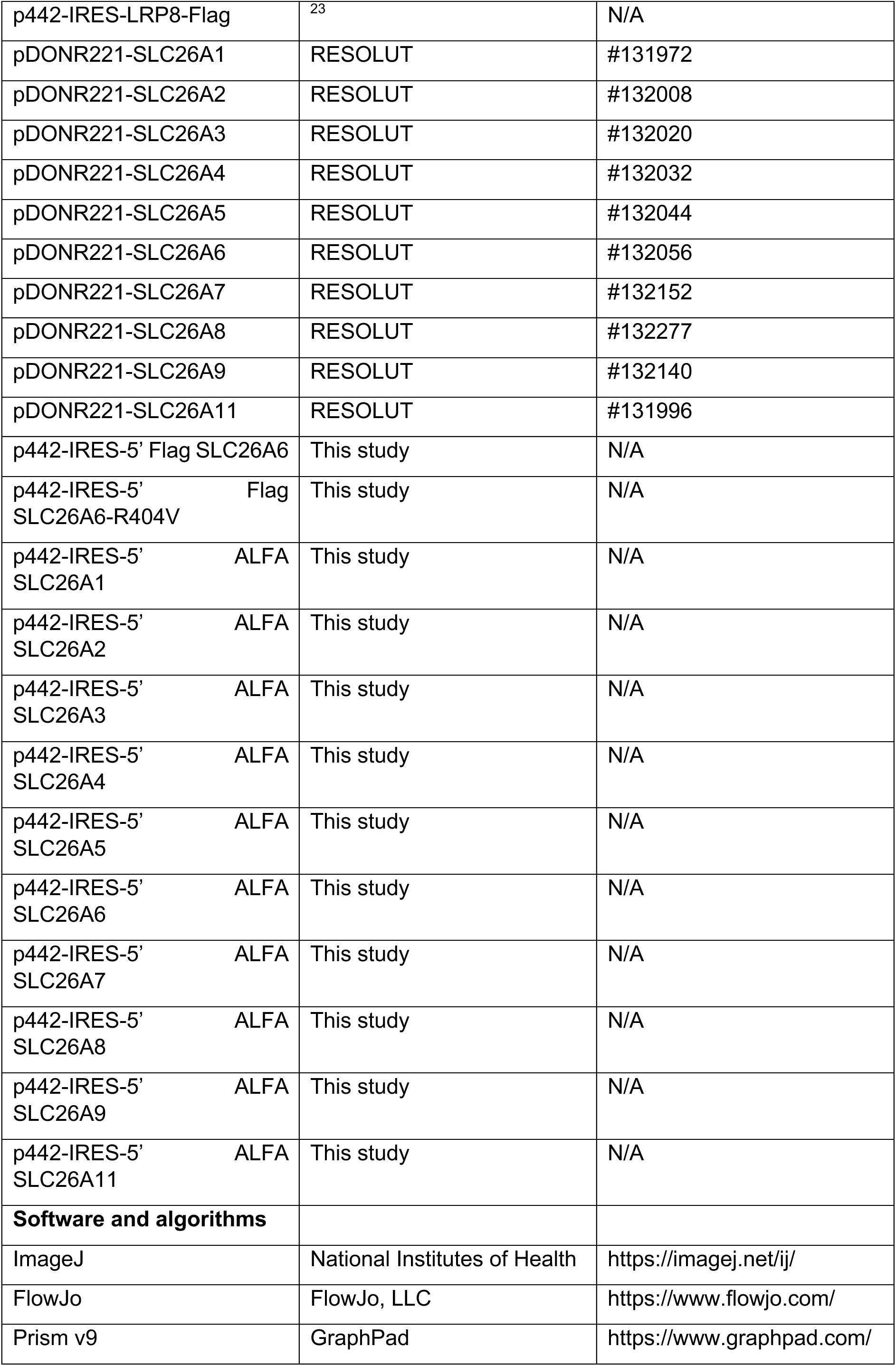
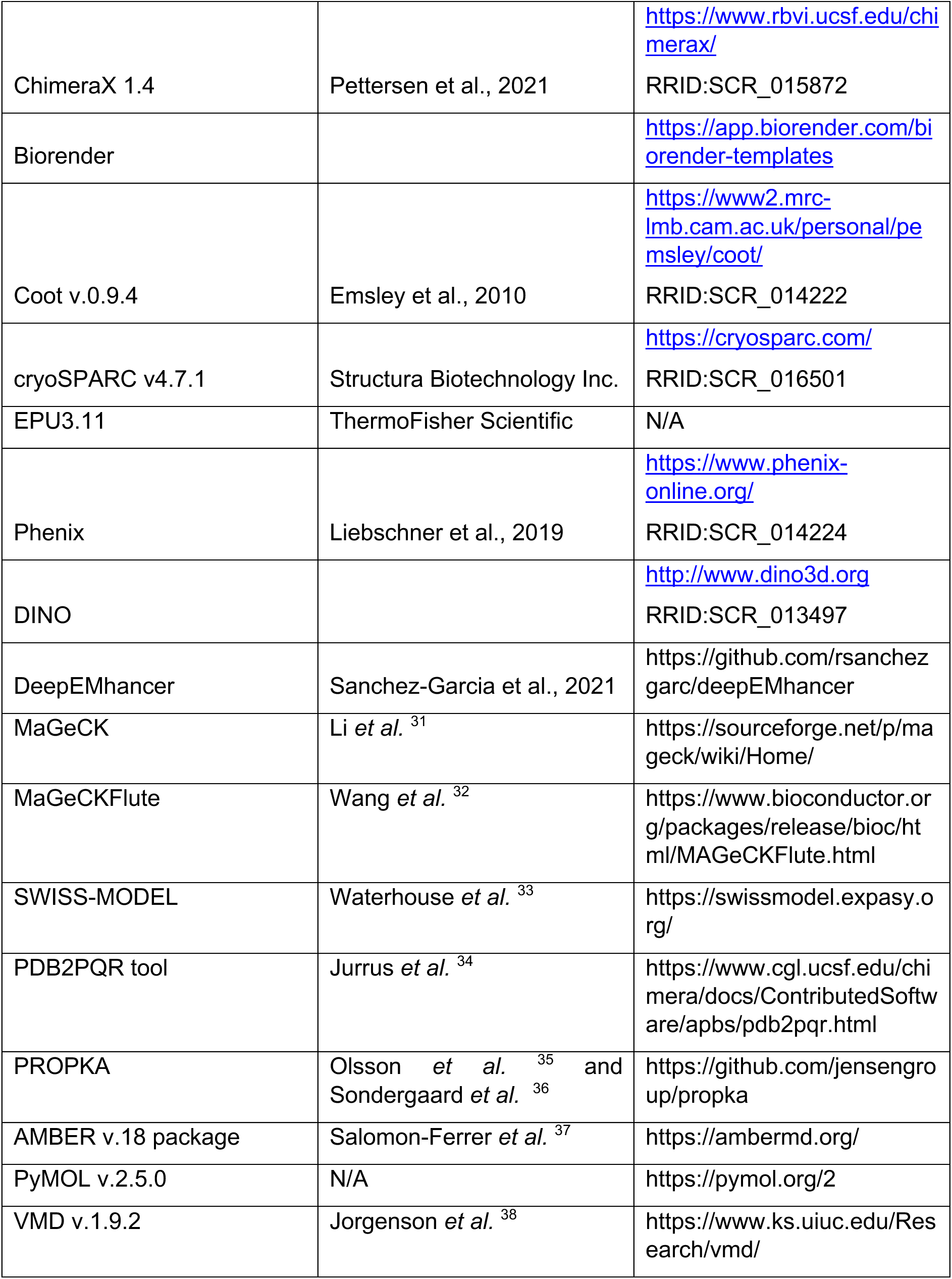

| **REAGENT or RESOURCE** | **SOURCE** | **IDENTIFIER** |
|---|---|---|
| **Antibodies** |  |  |
| β-actin | Sigma-Aldrich | A5441 |
| GPX4 | abcam | ab125066 |
| LRP8 | abcam | ab108208 |
| SELENON | Santa Cruz Biotechnology | sc-365824 |
| Flag | Sigma-Aldrich | F3165 |
| VINCULIN | Santa Cruz Biotechnology | sc-73614 |
| SLC26A6 | NovusBiologicals | NBP2-47273 |
| FluoTag®-X2 anti-ALFA | NanoTag Biotechnologies | N1502-At488-L |
| GPX1 | Cell Signalling | 3286 |
| SELENOS | Santa Cruz Biotechnology | sc-365498 |
| SLC7A11 | developed in Munich | antibody raised against a N-terminal peptide of SLC7A11, clone 15C2-111 |
| anti-mouse IgG, HRP-linked Antibody | Cell Signaling | 7076 |
| Anti-rabbit IgG, HRP-linked Antibody | Cell Signaling | 7074 |
| Anti-rat IgG, HRP-linked Antibody | Cell Signaling | 7077 |
| **Bacterial and virus strains** |  |  |
| NEB® 5-alpha Competent *E. coli* | New England Biolabs | C2987H |
| DH10Bac™ *E. coli* competent cells | Thermo Fischer |  |
| **Chemicals, peptides, and recombinant proteins** |  |  |
| Erastin | Selleckchem | S724203 |
| RSL3 | Sigma | SML2234-5mg |
| ML210 | Sigma | SML0521-5mg |
| BSO | Sigma | B2515-500MG |
| Fin56 | Sigma | SML1740-5mg |
| L-Selenocystine | Sigma | 545996-250MG |
| Lipoxstatin-1 | Sigma | SML1414-25mg |
| Sodium sulfite | Sigma | 239321 |
| Sodium sulfate | Sigma | 238597 |
| Sodium selenite | Sigma | S5261-10G |
| Sodium oxalate | Sigma | 223433 |
| Seleno-L-cystine | Sigma | 545996 |
| L-Selenomethione | Sigma | S3132-100MG |
| DMEM, low glucose, GlutaMAX™ Supplement, pyruvate | Thermo Fisher | 10567014 |
| DMEM, high glucose, no glutamine, no methionine, no cystine | Thermo Fisher | 21013024 |
| Penicillin-Streptomycin (10,000 U/mL) | Thermo Fisher | 15140122 |
| Dulbecco’s Phosphate Buffered Saline | Sigma-aldrich | D8537-24X500ML |
| GlutaMAX<sup>TM</sup> | Thermo Fisher | 35050038 |
| Sodium Pyruvate | Thermo Fisher | 11360039 |
| L-Methionine | Sigma | M9625 |
| resazurin sodium salt | Sigma |  |
| yttrium standard solution | Merck Millipore | 119809 |
| 10X PBS | Sigma | D1408-500ML |
| 4-Hydroxytamoxifen | Selleckchem | S7827 |
| Protease inhibitor cocktail | Roche | 04693159001 |
| enhanced chemiluminescence substrate | BioRad | 107-5061 |
| scintillation solution, Ultima Gold MV | PerkinElmer | #6013159 |
| 14C-oxalate (specific activity 116 mCi/mmol) | ViTrax |  |
| HyClone HyCell TransFx-H medium | Cytiva | SH30939.02 |
| SF-4 Baculo Express ICM 500mL | Bioconcept | 9-00F38-I |
| TransIT®-Insect Transfection Reagent | mirusbio (MERCK) | MIR 6100 |
| SLC26A6 inhibitor (A6i) | ChemDiv | Y021-4539 |
| A6i.01 | ChemDiv | 4151-0534 |
| A6i.02 | ChemDiv | 4477-0210 |
| A6i.03 | ChemDiv | 7377-1459 |
| A6i.04 | ChemDiv | 8016-0556 |
| A6i.05 | ChemDiv | 8016-4802 |
| A6i.06 | ChemDiv | C301-5961 |
| A6i.07 | ChemDiv | Y020-5758 |
| A6i.08 | ChemDiv | Y020-6003 |
| A6i.09 | ChemDiv | Y020-6941 |
| A6i.10 | ChemDiv | Y020-6943 |
| A6i.11 | ChemDiv | Y020-6944 |
| A6i.12 | ChemDiv | Y021-1529 |
| A6i.13 | ChemDiv | Y021-4410 |
| 1-palmitoyl-2-oleoyl-sn-glycero-3-phosphoethanolamine (POPE) | Avanti Polar Lipids | 850757C |
| 1-palmitoyl-2-oleoyl-sn-glycero-3-phospho-(1'-rac-glycerol) (POPG) | Avanti Polar Lipids | 840457C |
| Triton X-100 | Sigma | T9284 |
| Pepstatin A | Axon lab | A2205.0100 |
| Pepstatin A | Axon lab | A2205.0100 |
| Phenylmethylsulfonyl fluoride (PMSF) | Sigma | PMSF-RO |
| Leupeptin | AppliChem | A2183,0100 |
| Benzamidine | Sigma | B6506 |
| Chloroform | Fluka | 25690 |
| Glyco-diosgenin (GDN) | Anatrace | GDN101 |
| Cholesteryl Hemisuccinate Tris Salt (CHS) | Anatrace | CH210 |
| n-Dodecyl-β-D-Maltopyranoside (DDM) | Anatrace | D310 |
| Diethyl ether | Sigma | 296082 |
| DNase I | AppliChem | A3778 |
| Glycerin, Glycerol 86% | Roth | 4043.3 |
| HCl | Merck Millipore | 1.00319.1000 |
| HEPES | Sigma | H3375 |
| ACMA | Thermo Fischer | A1324 |
| Sodium butyrate | Sigma | 303410 |
| CCCP | Sigma | C2759 |
| Phosphate buffered saline | Sigma | D8537 |
| Strep-Tactin<sup>®</sup> Superflow<sup>®</sup> high capacity resin | IBA | 2-1208-010 |
| D-desthiobiotin | Sigma | D1411 |
| Kolliphor P188 | Sigma | K4894 |
| L-glutamine | Sigma | G7513 |
| Penicillin-streptomycin | Sigma | P0781 |
| Fetal bovine serum | Sigma | F7524 |
| Calcium Chloride | Sigma | 223506 |
| Magnesium Chloride | Fluka | 63,065 |
| Potassium chloride | Sigma | 746346 |
| Sodium chloride | Sigma | 71380 |
| BODIPY 581/591 C11 | Thermo Fisher | D3861 |
| T4 DNA Ligase | New England Biolabs | M020L |
| BstBI | New England Biolabs | R0519S |
| XhoI | New England Biolabs | R0146S |
| BsmBI | New England Biolabs | R0739 |
| Midori Green Advance Safe DNA/RNA stain | NIPPON Genetics EUROPE | MG04 |
| Wizard® SV Gel and PCR Clean-up System | Promega | A9282 |
| **Critical commercial assays** |  |  |
| Monarch Plasmid Miniprep Kit | New England Biolabs | T1010L |
| Clone Express MultiS One Step Cloning Kit | Vazyme | VB-C113-02 |
| ClonExpress II One Step Cloning Kit | Vazyme | VB-C112-02 |
| X-tremeGENE HP DNA Transfection Reagent | Sigma-Aldrich | 6366546001 |
| Pierce BCA Protein Assay Kit | Thermo Fisher | 23225 |
| Dialysis tubing cellulose membrane | Sigma | D9527-100FT (average flat width 43 mm)<br>D9402-100FT (average flat width 76 mm) |
| **Deposited data** |  |  |
| Western blots and microscopy images | This study | DOI: Data will be deposited before publication |
| Mass spectrometry data | This study |  |
| CRISPR screening data | This study | NCBI Sequence Read Archive (SRA): |
| **Experimental models: Cell lines** |  |  |
| SK-N-DZ | ATCC | CRL-2149 |
| A375 | ATCC | CRL-1619 |
| HT1080 | ATCC | CCL-121 |
| A549 | ATCC | CRM-CCL-185 |
| NICH1975 | ATCC | CRL-5908 |
| HEK293T | ATCC | CRL-3216 |
| UACC62 | ATCC |  |
| LOXIMVI | ATCC |  |
| NMB | DSMZ | ACC 657 |
| HepG2 | ATCC | HB-8065 |
| Mel-JUSO | Kindly provided by Alexander Roesch, University Hospital Essen |  |
| MeWo | Kindly provided by Alexander Roesch, University Hospital Essen |  |
| IGR37 |  |  |
| RPMI-7951 | ATCC | HTB-66 |
| UACC257 |  |  |
| SK-MEL-2 | ATCC | HTB-68 |
| SK-MEL-28 | ATCC | HTB-72 |
| SK-MEL-5 | ATCC | HTB-70 |
| SK-N-AS | ATCC | CRL-2137 |
| SK-N-SH | ATCC | HTB-11 |
| NGP |  |  |
| SH-SY5Y | ATCC | CRL-2266 |
| Huh7 |  |  |
| NCI-H520 | ATCC | HTB-182 |
| MDA-MB-231 | ATCC | CRM-HTB-26 |
| HEK293S GnTi- | ATCC | CRL-3022 |
| Sf9 | Thermo Fischer |  |
| 4-hydroxytamoxifen (TAM)-inducible Gpx4−/− murine immortalized fibroblasts | Kindly provided by Dr. Marcus Conrad, Helmholtz Munich | Pfa1 |
| **Experimental models: Organisms/strains** |  |  |
| NOD.Cg-Prkdc<sup>scid</sup> Il2rg<sup>tm1Wjl</sup>/SzJ (NSG) mice | JANVIER LABS | N/A |
| **Oligonucleotides** |  |  |
| See Table S1 | This study | Table S1 |
| **Recombinant DNA** |  |  |
| pMDLg/pRRE | Addgene | #12251 |
| pRSV-Rev | Addgene | #12253 |
| pCMV-VSV-G | Addgene | #8454 |
| pLenti-CRISPR-V2 | Addgene | #98293 and #98290 |
| Mammalian expression vector with C-terminal 3C cleavage site, venus fluorescent tag, myc tag, SBP tag. | Dutzler laboratory | N/A |
| pEZT-BM | Hibbs laboratory (addgene database) | 74099 |
| FX-compatible<br>pEZT-BM | Cristina Paulino lab<br>University of Heidelberg |  |
| Recombinant Protein<br>HRV 3C protease | Expressed (pET_3C) and purified in Dutzler laboratory | N/A |
| p442-IRES-Mock | Kindly provided by Dr. Marcus Conrad, Helmholtz Munich |  |
| SLC-focused knock-out CRISPR library | a kind gift from Giulio Superti-Furga | Addgene #90294 |
| p442-IRES-Mock | This study | N/A |
| p442-IRES-LRP8-Flag | <sup>23</sup> | N/A |
| pDONR221-SLC26A1 | RESOLUT | #131972 |
| pDONR221-SLC26A2 | RESOLUT | #132008 |
| pDONR221-SLC26A3 | RESOLUT | #132020 |
| pDONR221-SLC26A4 | RESOLUT | #132032 |
| pDONR221-SLC26A5 | RESOLUT | #132044 |
| pDONR221-SLC26A6 | RESOLUT | #132056 |
| pDONR221-SLC26A7 | RESOLUT | #132152 |
| pDONR221-SLC26A8 | RESOLUT | #132277 |
| pDONR221-SLC26A9 | RESOLUT | #132140 |
| pDONR221-SLC26A11 | RESOLUT | #131996 |
| p442-IRES-5’ Flag SLC26A6 | This study | N/A |
| p442-IRES-5’ Flag SLC26A6-R404V | This study | N/A |
| p442-IRES-5’ ALFA SLC26A1 | This study | N/A |
| p442-IRES-5’ ALFA SLC26A2 | This study | N/A |
| p442-IRES-5’ ALFA SLC26A3 | This study | N/A |
| p442-IRES-5’ ALFA SLC26A4 | This study | N/A |
| p442-IRES-5’ ALFA SLC26A5 | This study | N/A |
| p442-IRES-5’ ALFA SLC26A6 | This study | N/A |
| p442-IRES-5’ ALFA SLC26A7 | This study | N/A |
| p442-IRES-5’ ALFA SLC26A8 | This study | N/A |
| p442-IRES-5’ ALFA SLC26A9 | This study | N/A |
| p442-IRES-5’ ALFA SLC26A11 | This study | N/A |
| **Software and algorithms** |  |  |
| ImageJ | National Institutes of Health | https://imagej.net/ij/ |
| FlowJo | FlowJo, LLC | https://www.flowjo.com/ |
| Prism v9 | GraphPad | https://www.graphpad.com/ |
| ChimeraX 1.4 | Pettersen et al., 2021 | https://www.rbvi.ucsf.edu/chimerax/<br>RRID:SCR_015872 |
| Biorender |  | https://app.biorender.com/biorender-templates |
| Coot v.0.9.4 | Emsley et al., 2010 | https://www2.mrc-lmb.cam.ac.uk/personal/pemsley/coot/<br>RRID:SCR_014222 |
| cryoSPARC v4.7.1 | Structura Biotechnology Inc. | https://cryosparc.com/<br>RRID:SCR_016501 |
| EPU3.11 | ThermoFisher Scientific | N/A |
| Phenix | Liebschner et al., 2019 | https://www.phenix-online.org/<br>RRID:SCR_014224 |
| DINO |  | http://www.dino3d.org<br>RRID:SCR_013497 |
| DeepEMhancer | Sanchez-Garcia et al., 2021 | https://github.com/rsanchezgarc/deepEMhancer |
| MaGeCK | Li *et al.* <sup>31</sup> | https://sourceforge.net/p/mageck/wiki/Home/ |
| MaGeCKFlute | Wang *et al.* <sup>32</sup> | https://www.bioconductor.org/packages/release/bioc/html/MAGeCKFlute.html |
| SWISS-MODEL | Waterhouse *et al.* <sup>33</sup> | https://swissmodel.expasy.org/ |
| PDB2PQR tool | Jurrus *et al.* <sup>34</sup> | https://www.cgl.ucsf.edu/chimera/docs/ContributedSoftware/apbs/pdb2pqr.html |
| PROPKA | Olsson *et al.* <sup>35</sup> and Sondergaard *et al.* <sup>36</sup> | https://github.com/jensengroup/propka |
| AMBER v.18 package | Salomon-Ferrer *et al.* <sup>37</sup> | https://ambermd.org/ |
| PyMOL v.2.5.0 | N/A | https://pymol.org/2 |
| VMD v.1.9.2 | Jorgenson *et al.* <sup>38</sup> | https://www.ks.uiuc.edu/Research/vmd/ |

### Cell culture

Cells were maintained in Dulbecco’s Modified Eagle Medium (DMEM) (Gibco), supplemented with 10% FBS (Gibco) and 100U/mL penicillin-streptomycin at 37°C with 5% CO_2_. SK-N-DZ cells deficient for *LRP8* were kept in a complete medium supplemented with 500 nM Lip1. Cell lines were passaged by trypsinization when they reached 80–90% confluency. Cells were regularly tested for mycoplasma using MycoBlue Mycoplasma Detector kit (Vazyme Biotech, VB-D101-02).

### SLC-focused CRISPR knockout screen

A549 cells were transduced with SLC-focused knock-out CRISPR library at MOI of 0.3. Nine million cells were transduced to achieve > 1000 cells per sgRNA for 1000X coverage. After selection with 1 μg/mL puromycin and recovery, cells were split into two conditions of controls or treated with IC90 value of selenite (3 μM) for 3 day and maintained at the sgRNA coverage of > 1000X coverage. Cell pellets were harvested after 2 weeks of passaging, followed by genomic DNA purification (QIAGEN-DNeasy) and sequenced as a multiplexed pool on a Nextseq500. The mapping of raw sequencing reads to the reference library and computing of enrichment scores, and p-values were processed using the MaGeCK pipeline ^31^. The MaGeCKFlute was used to identify gene hits and associated pathways ^32^.

### Molecular cloning

cDNA fragments for SLC26 family were gifts from RESOLUTE Consortium & Giulio Superti-Furga as pDonr221 plasmids. These cDNAs were cloned into linearized lentiviral vector p442-IRES (kindly provided by Dr. Marcus Conrad) using Clone Express MultiS One Step Cloning Kit (Vazyme, # VB-C113-02).

Individual sgRNAs were cloned into the pLenti-CRISPR-V2 (Addgene #98293 and #98290) lentivectors for CRISPR Knock-out (CRISPR-KO) experiments via restriction digest of the respective lentivector with BsmBI (NEB, #R0739) and T4 ligase (NEB, M020L). Table S1 shows the oligonucleotides (synthesized by Eurofins Genomics or Microsynth) with the sgRNA sequences and complementary overhangs, which were annealed and inserted into the respective lentiviral delivery vectors. PCR products were ligated via Clone Express MultiS One Step Cloning Kit with linearized p442 vector.

**Table S2.**
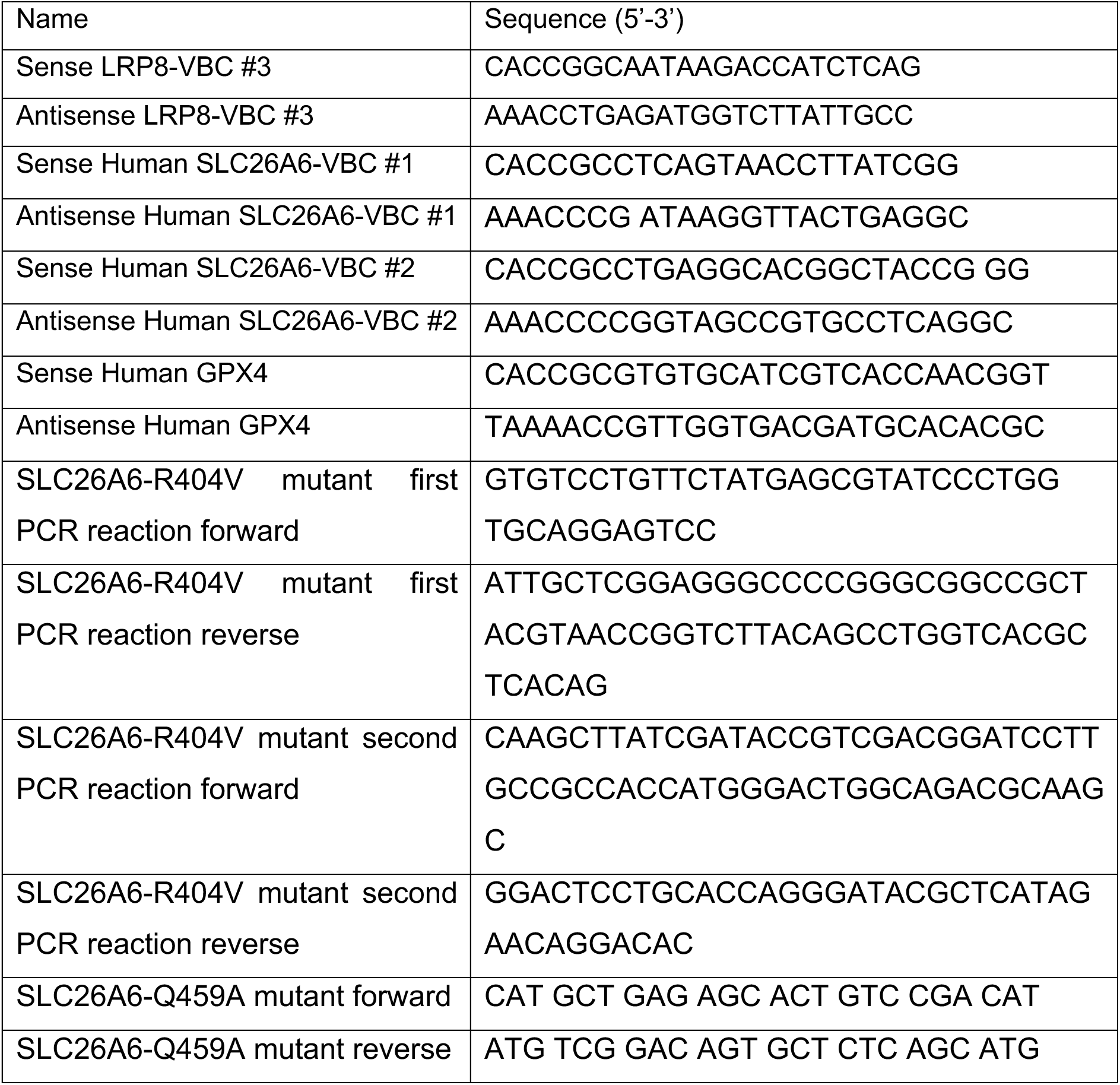
Oligonucleotides used for cloning individual gRNAs or cloning constructs. Related to the STAR Methods.

### Lentivirus production and transduction

Lentivirus was produced with a third-generation packaging system and X-tremeGENE HP DNA Transfection Reagent (Sigma-Aldrich, 6366546001). HEK293T cells were transfected with the lentiviral transfer plasmids, two packaging plasmids (pMDLg/pRRE, Addgene #12251 and pRSV-Rev, Addgene #12253), and the envelope expressing plasmid (pCMV-VSV-G, Addgene #8454). Viral supernatant was harvested 48 h post-transfection filtered through a 0.45 um filters and added to the target cells with 8 ug/mL protamine sulphate. Selection started 48 hours post transduction in the presence of antibiotics (puromycin 1 ug/mL, blasticidin 10 ug/mL, hygromycin B Gold 20 ug/mL, neomycin 100 ug/mL). All experimental procedures for lentivirus production and transduction were performed in a biosafety level 2 laboratory.

### Construct generation for cryo electron microscopy

The open reading frame of human SLC26A6 (isoform 3, GenBank accession number: BC017697) with a C-terminal venus YFP and streptavidin-binding peptide (SBP) tag was initially cloned into a pcDNA3.1 vector (Invitrogen) that had been modified to support FX cloning ^39^. For BacMam-mediated expression, the construct was subcloned into a modified version of pEZT-BM (Addgene plasmid # 74099) that was engineered to support FX cloning and to introduce a 3C protease cleavage site upstream of the Venus-SBP tag (generously provided by Cristina Paulino). The final expression construct encodes SLC26A6 containing a C-terminal fusion of 3C-venus-SBP with an additional serine at the N-terminus and alanine at the C-terminus resulting from the FX recombination system. The expression plasmid was transformed into DH10Bac™ *E. coli* competent cells, where Tn7 transposition was used to insert the construct into the bacmid genome. Positive colonies were selected and recombinant bacmid DNA was purified for transfection following the standard procedure for BacMam construct generation ^40^.

### Cell viability assays

The impact of chemical compounds or gene expression on cell viability was assessed using the Alamar Blue assay. To determine the cell viability, 3,000-4000 cells were seeded in a 96-well plate with standard medium (DMEM, 10% FBS + 1% penicillin/streptomycin) or cystine-free medium (DMEM with high glucose, no glutamine, no methionine, no cystine (Thermo Fischer Scientific, 21013024) supplemented with 1X GlutaMAX^TM^ (Thermo Fischer Scientific, 35050038), 1X Sodium Pyruvate (Thermo Fischer Scientific, 11360039), 0.2mM L-Methionine (Sigma-Aldrich, M9625) and 500 nM Lip1) 4 hours before the treatment. Cellular viability measurements were assessed after 48h of treatment unless otherwise stated. Cell viability after GPX4 inhibitor treatments was assessed after 72 h. Alamar blue stock solution was prepared by dissolving 1 g resazurin sodium salt (Sigma) in 100 mL sterile PBS and sterile filtrated through a 0.22 µm filter. Working solutions were prepared by adding 80 μl from the stock solution into 10 mL growth medium. After 2-3h incubation, viability was assessed by measuring the fluorescence using a 540±35 excitation filter and a 590±20 emission on a Spark® microplate reader (Tecan).

### Preparation of dialysed FBS

Dialysis tubing cellulose membrane (Sigma-Aldrich, D9527-100FT [avg. flat width 43 mm] or D9402-100FT [avg. flat width 76 mm]) was cut to fit the size of beaker. The membrane was soaked in distilled water for few minutes, and one end of the tube was sealed. FBS was transferred to the tube and the other end of the tube was sealed tightly. The serum was dialyzed against 1X PBS (prepared from 10X PBS, Sigma-Aldrich, D1408-500ML) at 1:10 (v/v) ratio in a beaker with magnetic stirring at 4°C for at least 5-6 hours or overnight. PBS was replaced and this step was repeated five times. Dialyzed FBS was sterilized using 0.2 µM filter.

### Immunoblotting

Cells or tissue samples were lysed in RIPA buffer (20mM Tris-HCl, 150mM NaCl, 1mM EDTA, 1mM EGTA, 1% NP-40, 1% deoxycholate) supplemented with the 1X protease inhibitor cocktail (Roche, 04693159001) and froze overnight to rupture the plasma membrane. After centrifuging the lysate for 30 min at 4°C and 18000g, supernatants were collected. Protein concentrations were measured using Pierce BCA Protein Assay Kit (Thermo Scientific, 23227) with bovine serum albumin as protein standard. 12% SDS-polyacrylamide gel electrophoresis was used to separate proteins followed by electro-transfer to polyvinylidene difluoride (PVDF) membranes. Membranes were blocked in 5% non-fat milk in Tris-buffered saline with 0.1 % Tween 20 (TBS-T) for 1 h at room temperature. Primary antibodies were diluted according to the manufacturer’s suggestion in a 3% bovine serum albumin (BSA) or 5% non-fat milk in TBS-T solution and incubated with the membrane overnight at 4°C. After 3 times washes in TBS-T, membranes were incubated with 1:3000 dilution of the secondary antibodies (Cell Signalling Technology, anti-rabbit IgG-HRP linked (#7074), anti-mouse IgG-HRP linked (#7076), or anti-rat IgG-HRP linked (#7077)) at RT for 2 hours. Proteins were detected using enhanced chemiluminescence substrate (BioRad, 107-5061). Images were acquired using detection systems Amersham ImageQuant 800 (Cytiva).

### Immunofluorescence staining

Cells expressing the ALFA tag constructs were seeded on glass coverslips in 6-well plates. On the next day, cells were washed with PBS and fixed in 4 % paraformaldehyde (Sigma) for 30 min, quenched in PBS containing 0.1 M glycine for 10 min, and permeabilized and blocked for 15 min in PBS containing 10 % normal goat serum and 0.1 % Triton X-100. Then, coverslips were incubated with FluoTag®-X2 anti-ALFA conjugated with Atto488 diluted 1:500 in 3 % normal goat serum and 0.1 % Triton X-100 for an hour. DNA was counterstained by incubating with DAPI (Thermo Fischer, 1:10000 in PBS) for 10 min, followed by two more PBS washes, 10 min each. The coverslips were then mounted to a microscopic slide using DABCO (Sigma). Confocal images were obtained using a Leica SP8 microscope with a HCX PL APO CS 63x 1.4 NA oil objective. Images were colorized, and the intensity was adjusted in ImageJ.

### Total reflection X-ray fluorescence (TXRF) analysis

Cells were seeded into 10cm dishes until they reached 90% confluency and then were treated with 50 µM sodium selenite in cystine free medium and 0.5 µM liproxstatin1 for 6 hours. Then the cells were collected via trypsinization as pellets, washed twice with PBS and split into 2 tubes and stored at −20°C. Cell pellets were lysed with RIPA buffer containing protease inhibitor as described earlier ^41^. Selenium measurements were performed using TXRF spectrometer (S4 T-STAR, Bruker Nano GmbH, Berlin, Germany). As internal standard 1mg/L yttrium (Merck Millipore, 119809) was used, and samples were loaded to siliconized quartz glass carriers (Serva, 35130.01) and dried at 40°C. Samples were measured in duplicates for 1000 s and selenium levels were normalized to protein concentration determined by Bradford analysis.

### Competition assay

Growth dynamics and competition of cells were analysed using the IncuCyte bioimaging platform (Sartorius). For this, A549 nontarget and sgSLC26A6 cells were transduced with mCherry-NLS (nuclear localization signal) and eGFP-NLS respectively and were seeded in 96-well plates (2000 cells per well for each cell line). Treatment with the indicated concentrations of sodium selenite started 4 hours later to allow cell attachment. Starting from time zero of treatment, two images per well were captured every 6 hours, averaged, and analyzed. Data were collected as count of red or green signals per total number of cells per image.

### Small molecule library screen

HT1080 cells overexpressing hSLC26A6 were seeded on 384-well plates (1,000 cells per well) in cystine-free medium (DMEM with high glucose, no glutamine, no methionine, no cystine) supplemented with 10% FBS, penicillin-streptomycin, L-glutamine (0.2 mM), sodium pyruvate (1 mM), 0.2 mM L-methionine (0.2 mM) and Lip1 (500 nM). After cell attachment, individual compounds from a small-molecule compound library (ChemDiv Library and ChemDiv Protein-Protein Interaction Library, total of 20000 compounds) were added at a final concentration of 10 μM, followed by the addition of sodium selenite (50 μM). Cell viability was assessed 24 h after selenite treatment using a resazurin reduction assay. Compounds that exhibited a rescue effect in the primary screening were further evaluated at concentrations ranging from 0.1 to 10 μM under the same conditions to validate their efficacy.

### Oxalate uptake measurement

To assess the capacity of cells for oxalate uptake, 200,000 cells per well were seeded in a 24-well plate, the uptake assay was performed 48 hours after seeding. The following conditions were tested in triplicates: chloride-containing buffer (5 mM glucose, 130 mM KCl, and 20 mM HEPES, pH 7.4), chloride-free buffer (5 mM glucose, 130 mM K-gluconate, 20 mM HEPES, pH 7.4), and chloride-free buffer + A6 inhibitor. The inhibitor was used at a final concentration of 5 µM and was added 30 minutes prior to measurement and kept during the incubation with ^14^C-oxalate. Before the measurement, cells were washed three times with chloride-free buffer. Cells were incubated with 5 μM ^14^C-oxalate (specific activity 116 mCi/mmol, ViTrax, Placentia, CA, US) for 4 minutes. Afterwards, the radioactive solution was aspirated, and the cells were washed three times with ice-cold chloride-free buffer. Cells were lysed with 200 μL of 0.2 N NaOH for 5 minutes, followed by 200 μL of 0.2 N HCl. Cell lysates were mixed with 3 mL of scintillation solution (#6013159, Ultima Gold MV, PerkinElmer) and counted using a Tri-Carb 4810TR scintillation counter (PerkinElmer, Waltham, MA, USA). A separate plate was prepared for cells in parallel for protein isolation and used for normalization of the data to total protein content.

### *In vivo* orthotopic mouse experiments

Following German legal regulations, the Regierungspraesidium Karlsruhe, the state’s governmental review board, approved all studies involving mice and experimental protocols under the authorization number G-176/19. The studies were carried out in accordance with the guidelines of the German Cancer Centre Institute. NSG mouse strains were used in the study. Experiments were conducted using female mice aged 3-4 months. Mice were kept in separately ventilated cages under temperature and humidity control with *ad libitum* access to food. The cages were provided with an improved habitat, including bedding material. To generate orthotopic mouse models for neuroblastoma, 2 × 10^5^ of luciferase-expressing SK-N-DZ with *LRP8^WT^/SLC26A6^WT^*, *LRP8^WT^/SLC26A6^OE^*, *LRP8^KO^/SLC26A6^WT^*, and *LRP8^KO^/SLC26A6^OE^*cells were transplanted into the right adrenal gland after the surgical site was prepared. Cells were resuspended in a 1:1 (vol/vol) mix of growth factor-reduced matrigel (Corning) and PBS. A total of 20 μL of this cell suspension was injected into the right adrenal gland of anesthetized mice. Following tumour cell transplantation, we used an IVIS Spectrum Xenogen device (Calliper Life Sciences) to monitor the mice for bioluminescent signals indicating tumour growth. One week after injecting the cells, we noticed a clear signal from the tumours. Animal health was evaluated on a regular basis, and the mice were sacrificed as soon as they met the abortion criteria outlined in the procedure. The sample size was calculated with the help of a biostatistician using R version 3.4.0. Power analysis assumptions were as follows: α error = 5%, β error = 20%. Mice were randomly assigned to treatment groups prior to treatment. Animals were excluded from further evaluations if they had to be sacrificed before reaching the predetermined endpoint (due to weight loss or other termination circumstances). The animal groups were anonymized during *in vivo* experiments and follow-up assessments.

### Protein expression and purification

Purified bacmid DNA was transfected into Sf9 insect cells using TransIT-Insect Transfection Reagent. After 72 hours, the initial P0 virus stock was harvested from the culture supernatant through centrifugation at 500 g for 15 min and filtering through a 0.22 nm filter. This stock was stored with 2% FBS (fetal bovine serum) and then used to infect fresh Sf9 cells (1.2 10^6^ ml^-1^) in two sequential rounds (P1 and P2) to amplify the virus. The resulting amplified BacMam virus was subsequently used for transduction of mammalian cells for transient protein expression. To this end HEK293S GnTI^-^ (ATCC CRL-3022) cells were grown in suspension and maintained in HyClone TransFx-H (GE Healthcare) media supplemented with 1% FBS, 1% Kolliphor P188, 100 U ml^−1^ penicillin/streptomycin and 2 mM Glutamine. Cells were incubated at 37 °C and 5% CO_2_ in TubeSpin Bioreactor 600 vessels (TPP) and shaken at 185 rpm.

For protein expression, cells were split 24 h prior to viral transfection from a cell density of 1-1.5 10^6^ ml^-1^ to 0.4-0.6 10^6^ ml^-1^. The following day, amplified P2 virus carrying the expression construct was mixed with the cell culture in a ratio of 1:10. After another 24 h, cells were provided with sodium butyrate to a final concentration of 10 mM, the growth temperature was dropped to 30 °C and cells were incubated for another 48 h. Cells were then harvested and flash-frozen in liquid N_2_ prior to storage at −80 °C.

For protein purification, all steps were performed at 4 °C. Cell pellets were thawed and mixed with extraction buffer (25 mM HEPES, pH 7.4, 200 mM NaCl, 5% glycerol, 2 mM CaCl_2_, 2 mM MgCl_2_, 1.7 mM n-Dodecyl-β-D-Maltopyranoside (DDM), 0.27 mM Cholesteryl hemisuccinate (CHS)), 1 mM benzamidine, 10 µM leupeptin, 1 µM pepstatin A, 100 µM PMSF), 10 µg/mL DNAse. For lysis cells were incubated for 2 h at 4 °C under gentle agitation prior to ultracentrifugation (85,000 g for 30 min at 4 °C) to remove insoluble material. The lysate was filtered (5 µm filter, Sartorius) prior to binding to Streptactin Superflow high-capacity resin (IBA Lifesciences) equilibrated in wash buffer (25 mM HEPES, pH 7.4, 200 mM NaCl, 5% glycerol, 90 μM GDN (glyco-diosgenin)) for 2 h. The Streptactin resin was washed with 40 column volumes (CV) of wash buffer and SBP-tagged protein was eluted using wash buffer supplemented with 10 mM desthiobiotin (Sigma). The eluate was concentrated to 500 µl using a 100 kDa MWCO concentrator (amicon) pre-equilibrated in wash buffer. C-terminal purification and detection tags were cleaved by 3C protease at a 2:1 protease:protein mass ratio. Prior to size-exclusion chromatography, samples were passed through a 0.22 µm filter (Millipore) and injected on a Superose 6 10/300 column (GE Healthcare) in SEC buffer (10 mM HEPES, pH 7.4, 200 mM NaCl, 0.02% GDN). Protein-containing peak samples were collected and used either for cryo-EM sample preparation at a concentration of 3 mg ml^-1^ or for liposome reconstitution.

For the preparation of SLC26A6 in the absence of Cl, a similar purification approach was used except that the protein was eluted from the streptactin resin using buffer, containing 25 mM HEPES, pH 7.4, 200 mM sodium gluconate, 90 μM GDN and 10 mM desthiobiotin. During size exclusion chromatography, the buffer contained 10 mM HEPES, pH 7.4, 200 mM sodium gluconate and 0.02% GDN. Protein containing peaks were concentrated to 3.3 mg ml^-1^ and sodium selenite was added to a final concentration of 50 mM during an 1h incubation before vitrification on grids.

### Liposome Reconstitution and Transport Experiments

Lipid preparation was performed on the same day as protein purification. POPE:POPG lipids (3:1 w/w) were pooled in a round-bottom flask and initially dried under N_2_ flow to form a lipid film. Lipids were further washed with diethyl ether and dried again under vacuum for 1 h prior to re-hydration with buffer (10 mM HEPES, 150 mM KCl, 7.4 pH) to 20 mg ml^-1^. The resulting suspension was gently sonicated and stirred for 2 h to generate a homogeneous mixture. Ten freeze-thaw cycles were performed to generate multilamellar liposomes before extrusion 29 times through x2 polycarbonate 400 nm filters to obtain unilamellar liposomes.

For protein incorporation, liposomes were diluted to 4 mg ml^-1^ and 10 µl aliquots of 10% Triton X-100 were added in order to destabilize liposomes and permit protein incorporation. After reaching a plateau of the light scattering measured at 540 nm, 4 additional aliquots of Triton X-100 were added. The number of additions required for destabilization did not vary between reconstitutions. After the formation of destabilized liposomes, purified SLC26A6 was added at a lipid-to-protein ratio (LPR) of 50:1. The subsequent mixture was gently rotated at RT for 20 min prior to the addition of 250 mg SM-2 Bio-Beads (Bio-Rad) per 5 ml of sample. The sample was then incubated for 30 min at RT prior to its incubation at 4 °C. Aliquots of Biobeads were added over a three-day period every 24 h. Biobeads were removed by filtration and the sample was used immediately for assay measurements. The successful incorporation of protein was confirmed by SDS-PAGE. For all reconstitutions, mock liposomes lacking purified protein were generated using the same protocol.

To characterize the electrogenic chloride/oxalate exchange properties of SLC26A6, proteoliposomes were prepared with an internal buffer concentration of 150 mM KCl, 10 mM HEPES, pH 7.4 and sonicated in 100 µl aliquots to form unilamellar vesicles prior to dilution into 9-amino-6-chloro-2-methoxyacridine (ACMA) assay buffer (2 µM ACMA, 9.4-150 mM Na_2_C_2_O_4_, 10 mM HEPES, pH 7.4) to a final concentration of 0.8 mg ml^-1^. 100 µl aliquots of proteoliposome suspensions were added to a 96-well plate and the ACMA fluorescence signal (excitation at 412 nm and emission at 482 nm) was recorded on a Tecan Spark Plate reader. After measuring the baseline during a period of 13 excitation/emission cycles (0.5 s per cycle), the protonophore carbonyl cyanide 3-chlorophenylhydrazone (CCCP) was added (to reach a final concentration of 300 nM) to dissipate the membrane potential and permit transporter-mediated anion-movement. The ACMA fluorescence decrease, as indicator of electrogenic Cl**^−^**/C_2_O_4_^2**−**^-exchange mediated H^+^ influx, was measured over 1000 s. Data were normalized to the corresponding value after CCCP addition. For the characterization of the inhibition of Cl**^−^**/C_2_O_4_^2**−**^ exchange upon addition of 26A6i03 (A6i), a similar method was used. A6i was added as 20 mM stock solution in DMSO (0.2 μm filtered). Proteoliposomes were diluted with assay buffer to 0.8 mg ml^-1^, and incubated with A6i (in a concentration range between 0.4 - 15 µM) for 30 min. Subsequently, ACMA fluorescence was measured from 100 µl aliquots over 1000 s after CCCP addition. Transport experiments were repeated five times from two independent reconstitutions for SLC26A6 WT and five times from one reconstitution for of the mutant Q459A. Activities were quantified as the slope of the mean ACMA decay in the time range between 120 and 320 sec after substrate addition in case of WT and between 220 and 400 sec for Q459A.

For the assay of electrogenic selenite/oxalate exchange by SLC26A6, a similar strategy was used. Proteoliposomes were diluted in sodium selenite buffer (150 mM, 10mM Hepes, pH 7.4) and subjected to five freeze-thaw cycles, followed by sonication. The liposomes were harvested by centrifugation at 215,000 g for 45 min and resuspended in assay buffer (2 µM ACMA, 150 mM Na_2_C_2_O_4_, 10 mM HEPES, pH 7.4). 100 µl aliquots of proteoliposome suspensions were added to a 96-well plate to record the ACMA fluorescence signal after the addition of CCCP, following the same steps as with 26A6i03. Transport experiments with selenite were repeated eight times from two independent reconstitutions and data was normalized to the corresponding value after CCCP addition. An equivalent procedure was used for the assay of sodium gluconate transport used as negative control.

### Cryo-EM Data Acquisition

For structure determination by cryo-EM, 2.5 µl samples of GDN-purified SLC26A6 that was incubated with A6i at a final concentration of 0.6 mM, were applied to glow-discharged holey carbon grids (Quantifoil R1.2/1.3 Au 200 mesh) at a concentration of 3 mg ml^-1^. Blotting of the sample (3-6 s) was performed at 4 °C at a relative humidity of 100% and grids were flash-frozen in liquid propane-ethane mix with a Vitrobot Mark IV (Thermo Fisher Scientific). A total of 13,000 images were collected on a 300 kV Titan Krios G3i using a 100 μm objective aperture. Data were collected in super-resolution mode using a post-column quantum energy filter (Gatan) with a 20-eV slit and a K3 Gatan direct detector. Data were acquired using EPU 3.11 with aberration-free image shift (AFIS) and a defocus range of −1.1 to −2.6 μm. The dataset was recorded using a pixel size of 0.651 Å/pixel (0.3225 Å/pixel in super-resolution mode) with a total exposure time of 1 s (36 individual frames) and a dose of 1.7 e^-^/Å^2^/ frame. The total electron dose for the specimen was 61.34 e^-^/Å^2^. The SLC26A6 selenite samples were prepared as described above. GDN-purified SLC26A6 at 3.3 mg ml^-1^ was incubated with 50 mM sodium selenite for 1 h, before freezing on glow-discharged holey carbon grids (Quantifoil R1.2/1.3 Au 200 mesh). Blotting conditions were identical to the A6i sample. A total of 98,000 images were collected on a 300 kV Titan Krios G3i, while the total electron dose was 64.3 e**^−^**/Å^2^. The significantly larger dataset size was a result to compensate for the particle aggregation and lower contrast observed after the addition of selenite, compared to data obtaine for the SLC26/Cl**^−^**/A6_i_ complex.

### Cryo-EM data processing

Cryo-EM data were processed using CryoSPARC v4.7.1 ^42^. All movies were subjected to patch motion correction. Following patch CTF estimation, high-quality micrographs were identified based on relative ice thickness, CTF resolution estimation and total full frame motion. For the SLC26A6/Cl**^−^**/A6_i_ complex 1,300,000 particles were picked following the generation of 2D templates for automated template picking. Subsequently, picked particles were binned 2x using a box-size of 360 pixels for 2D classification (pixel size 1.56 Å/pixel).

After multiple rounds of 2D classification for the exclusion of junk particles, the best classes were pooled and used for 3-D reconstruction. Multiple ab-initio jobs were performed with different number of classes in order to acquire the optimal particle alignment. Once the best class was determined, particles were re-extracted with an unbinned pixel size of 0.651 Å/pixel using a 448 pixel-sized box and subjected to non-uniform refinement. The refined map with a nominal resolution of 2.72 Å was further refined using the DeepEMhancer tool using the HighRes deep-learning model ^43^.

For the SLC26A6/selenite sample, the data was processed in CryoSPARC, following a similar iterative particle picking routine. The large size of the dataset led to extensive rounds of 2D classification and gradual improvement of 3D reconstructions. Box sizes, particle re-extraction and final refinement through DeemEMhancer were performed in the same way as for the SLC26A6/Cl**^−^**/A6_i_ complex.

### Cryo-EM model building and refinement

For both datasets, map interpretation was performed in Coot ^44^ using a model of SLC26A6 obtained from UniProt Database 8OPQ ^16^ as template. The model was iteratively improved by real space refinement in PHENIX ^45^ maintaining secondary structure constrains throughout. Coordinates and restraints of A6i were generated with Frog v2.14 ^46^ and PHENIX and its structure within the complex was refined with PHENIX. Figures were generated using ChimeraX ^47^ and DINO (www.dino3d.org).

### Quantification and statistical analysis

Statistical analyses for each experiment are indicated in the corresponding figure legends and p-values are given in the figures. Mean ± SEM are depicted in graphs, unless otherwise stated. The number of independent repeats is mentioned in corresponding figure legends. Graphs were generated using GraphPad Prism v10 (GraphPad Software) and schemes were generated at BioRender. For data obtained using the Incucyte imaging system, Incucyte 2023A Rev2 software was used for data analysis.

### Chemical characterization of A6i

**Figure S9:**
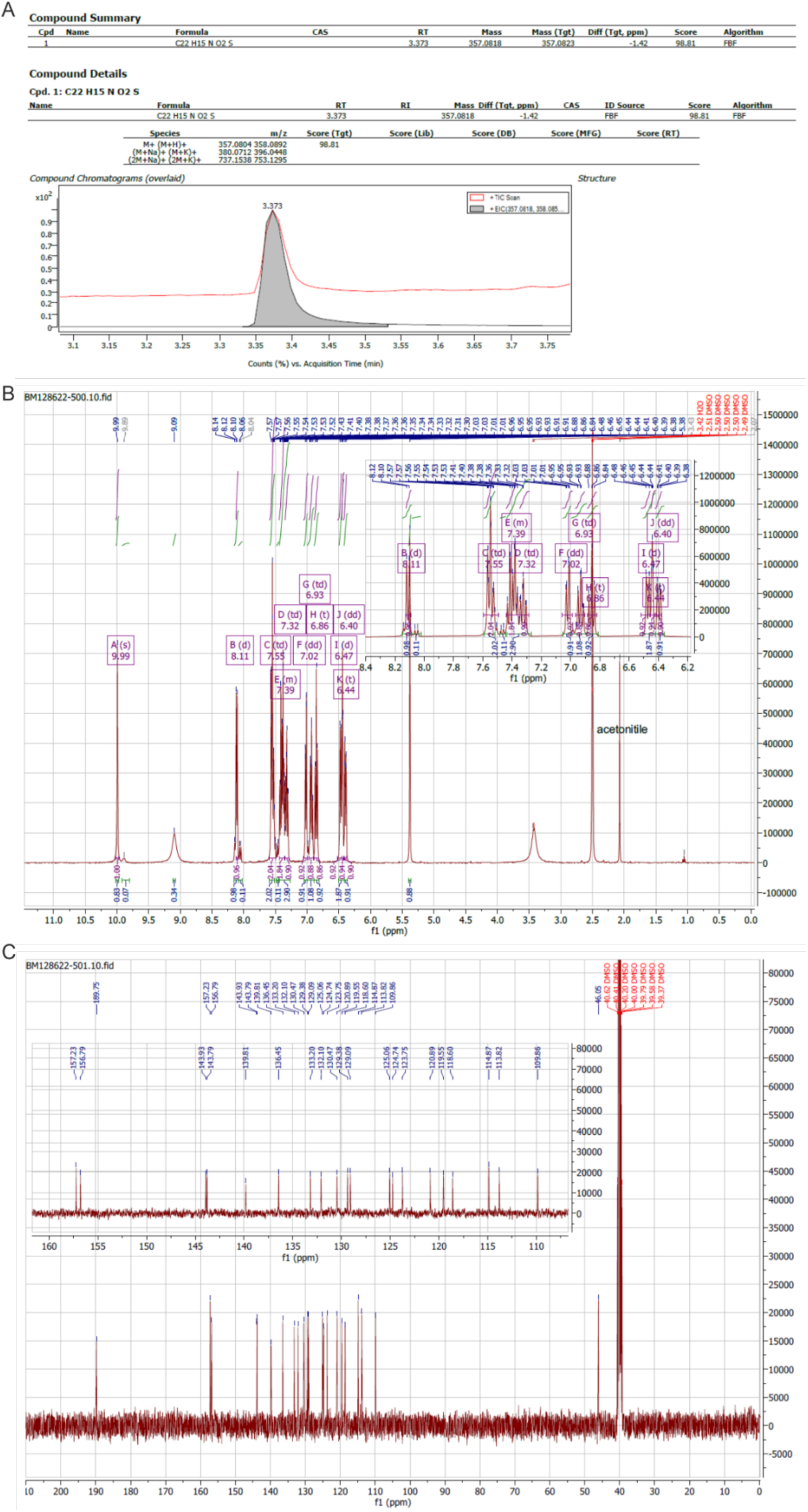
HRMS, ^1^H and ^13^C NMR analyses for A6i. **A)** HRMS **B)** ^1^H NMR **C)** ^13^C NMR results.

### Data, code, and materials availability

The coordinates Cryo-EM density map and the atomic model of the SLC26A6/selenite and the SLC26A6/Cl**^−^**/A6i complexes were deposited in the Protein Data Bank under accession codes 34YC and 9T11. The cryo-EM maps were deposited in the Electron Microscopy Data Bank under EMDB IDs EMD-100247 and EMD-55417.

This paper does not report any original code.

Any additional information required to reanalyze the data reported in this work paper is available from the lead contact upon request.

**Table S3.**
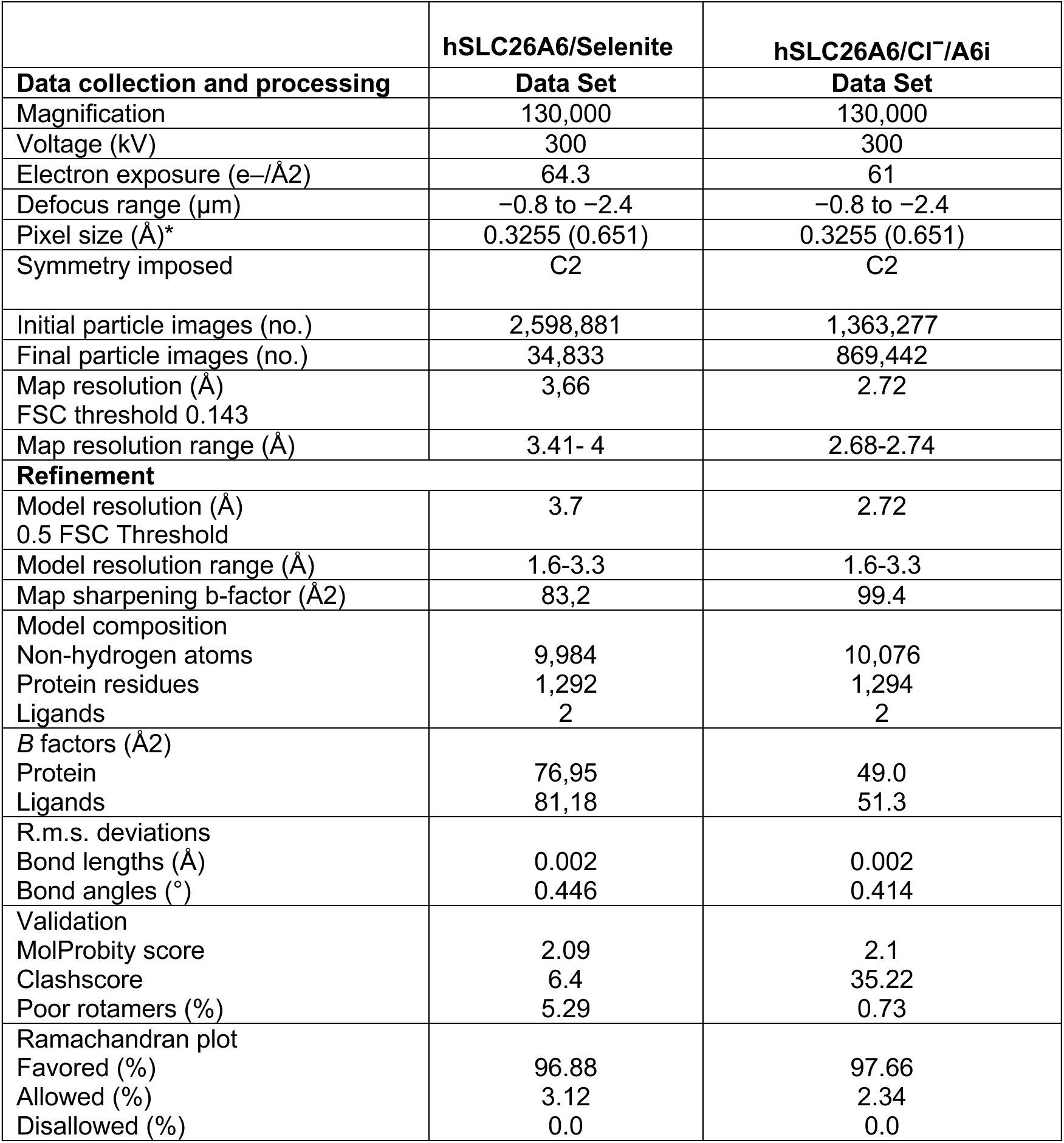
Cryo-EM data collection, refinement and validation statistics.

|  | **hSLC26A6/Selenite** | **hSLC26A6/Cl<sup>−</sup>/A6i** |
|---|---|---|
| **Data collection and processing** | **Data Set** | **Data Set** |
| Magnification | 130,000 | 130,000 |
| Voltage (kV) | 300 | 300 |
| Electron exposure (e–/Å2) | 64.3 | 61 |
| Defocus range (μm) | −0.8 to −2.4 | −0.8 to −2.4 |
| Pixel size (Å)* | 0.3255 (0.651) | 0.3255 (0.651) |
| Symmetry imposed | C2 | C2 |
| Initial particle images (no.) | 2,598,881 | 1,363,277 |
| Final particle images (no.) | 34,833 | 869,442 |
| Map resolution (Å)<br>FSC threshold 0.143 | 3,66 | 2.72 |
| Map resolution range (Å) | 3.41- 4 | 2.68-2.74 |
| **Refinement** |  |  |
| Model resolution (Å)<br>0.5 FSC Threshold | 3.7 | 2.72 |
| Model resolution range (Å) | 1.6-3.3 | 1.6-3.3 |
| Map sharpening b-factor (Å2) | 83,2 | 99.4 |
| Model composition<br>Non-hydrogen atoms<br>Protein residues<br>Ligands | 9,984<br>1,292<br>2 | 10,076<br>1,294<br>2 |
| *B* factors (Å2)<br>Protein<br>Ligands | 76,95<br>81,18 | 49.0<br>51.3 |
| R.m.s. deviations<br>Bond lengths (Å)<br>Bond angles (˚) | 0.002<br>0.446 | 0.002<br>0.414 |
| Validation<br>MolProbity score<br>Clashscore<br>Poor rotamers (%) | 2.09<br>6.4<br>5.29 | 2.1<br>35.22<br>0.73 |
| Ramachandran plot<br>Favored (%)<br>Allowed (%)<br>Disallowed (%) | 96.88<br>3.12<br>0.0 | 97.66<br>2.34<br>0.0 |

